# Live and let die: lysis time variability and resource limitation shape lytic bacteriophage fitness

**DOI:** 10.1101/2025.07.21.665848

**Authors:** Aaron Smith, Michael Hunter, Somenath Bakshi, Diana Fusco

## Abstract

Bacteriophages (phages) play a critical role in controlling bacterial populations, both in nature and as potential therapeutic agents. Their ability to replicate, compete against each other, and eradicate target cell populations is usually understood through a number of ‘life history parameters’, traditionally measured by population-level assays, which implicitly average the parameter’s value across a large number of infection events. Recent experiments suggest that bacteriophage life history parameters are subject to considerable heterogeneity, raising the question of whether experimental and modelling efforts that do not account for this variability may overlook important factors in phage’s behaviour, competitive fitness or therapeutic viability.

Here, using agent-based simulations, we investigate the importance of stochasticity in lysis time and burst size of lytic bacteriophages in two common laboratory competition experiments: serial passage of well-mixed populations and plaque expansion across a bacterial lawn. We find that a phage’s analytic growth rate in isolation can be a poor predictor of its fitness advantage in simulated competition experiments. Specifically, when lysis times are tightly distributed, we identify a novel effect we name “population resonance”, through which a bacteriophage can display a significant fitness advantage over a competitor with a much greater growth rate in isolation. Our simulations also show that both serial passage and plaque expansion reward variability in lysis time more than expected, by increasing the phage resilience when resources are scarce.

**Author Summary:** Bacteriophages (viruses that infect bacteria) can be described by a set of attributes called ‘life history parameters’. Historically, these parameters could only be measured on-average over large populations. Recent experimental advancements, however, have enabled their quantification at the single-cell-single-virus level, and have revealed that these traits are subject to inherent variability, even for genetically identical viruses infecting genetically identical cells. Here, we explore whether the degree of variability in one particular life history parameter, the lysis time (the time taken for a bacteriophage to infect and kill its host bacteria) might be subject to natural selection, by quantifying its effects on phage’s fitness in two simulated competition experiments: serial passage and plaque expansion. Surprisingly, we find that lysis time variability is advantageous in competition even when it reduces the phage’s growth rate in isolation. This finding opens the door to future computational and experimental work, as it demonstrates that mean values alone are not sufficient to describe or predict a bacteriophage efficacy, and that models that ignore this variability can overlook viable evolutionary strategies.

## 1 Introduction

Bacteriophages (phages) play a critical role in controlling bacterial populations both in nature, in which it is estimated that bacteriophages are responsible for 20 to 40% of all cell lysis events globally [1], and in medicine, where they provide a potential solution to the increasing spread of antimicrobial-resistance among human pathogens [2, 3]. In fact, bacteriophages have been successfully used as antimicrobials since Felix d’Herell’s experiments on rabbits in 1917 [4, 5], and recent trials have shown encouraging results [6, 7].

Bacteriophages can be described by a number of ‘life history parameters’, including adsorption rate (how readily the phage binds to target cells), lysis time (the time between adsorption to the cell and lysis), and burst size (the number of offspring released at cell death). The phenotypic assays typically used to measure these parameters were first developed several decades ago, and rely on population-level measurements, which inevitably average a parameter’s value across a large number of infection events [8–10]. These assays have very recently seen some innovation to accommodate cell-to-cell variability [11–13], but in experiments focussed on phage evolution, competition and optimality, measuring the population average life history parameters remains the popular approach [14–19].

In parallel to this trend, most computational and theoretical work on bacteriophages has traditionally employed ordinary or partial differential equation methods, modelling the progression from infection to lysis either as an explicit and deterministic delay, [20–23], as an exponential process [24, 25], or as a chain of intermediate steps connected by exponential processes [26]. These approaches assume that the distribution of lysis times will conform to a delta function, an exponential distribution, or an Erlang distribution, respectively. Some such work considers plasticity in viral parameters in response to the availability of nutrients to the bacterial host [27, 28]. Accounting for such extrinsic variation allows for a deeper understanding of phage population and evolutionary dynamics, but the intrinsic variation that exists even between clonal cells in identical conditions [29] (and therefore between infection outcome) remains unexplored.

Agent-based models (ABMs), in which bacteria and/or phages are treated as discrete objects with stochastic behaviour, are particularly well suited to investigate the role of noise in population dynamics. Specifically for phage populations, they have been successfully used to study the evolutionary consequences of a phage permitting or preventing superinfection [30], the physical volume occupied by bacteria in a growing biofilm [31, 32], the decay in heterozygosity at the front of an expanding plaque [33], and the possibility of a phage evolving towards symbiosis [34]. By construction, ABMs introduce a degree of variability, since the progression from each time step to the next is probabilistic, but few [30], to our knowledge, have incorporated explicit heterogeneity in lysis time or burst size.

In stark contrast to the general trend in the field summarized above, very recent experiments suggest that bacteriophages are subject to considerable heterogeneity in adsorption rate [16], lysis time [16, 35, 36] and burst size [12, 16, 37], the latter varying by more than an order of magnitude even in genetically identical cells in homogeneous conditions [37]. Since phages are totally dependent on the molecular machinery of their host, this variability likely arises from the considerable heterogeneity of the target bacteria’s internal metabolic state, via the availability of ribosomes [38–40], surface receptors [41, 42] and other critical molecules. Adsorption rate has also been shown to be influenced by environmental conditions such as pH and temperature directly [43]. Experimental and modelling efforts that do not account for such variability may, therefore, overlook critical factors that influence phage’s behaviour, competitive fitness or therapeutic viability [44].

While the ultimate source of variability in life history parameters likely arises from the host cells, experimental evidence suggests that a bacteriophage can modulate this variability, as different genetic backgrounds exhibit it at different levels [35, 36, 45]. We may therefore think of a “noisy phage” as a phage that possesses some genetic mechanism to amplify the biological noise of its host, and vice versa, a “non-noisy phage” as one suppressing it. One genetic mechanism which has been shown, for instance, to influence lysis time variability is stochastic expression of holins, proteins which create gaps in the host bacteria’s surface membrane [35, 36]. As long as these mechanisms are encoded in the phage genome, we can think of them as inheritable traits subject to selection.

In bacterial populations, variability in doubling time has been shown to result in a population growing faster than its mean doubling time would predict [46, 47], but whether this result holds in phage populations is not obvious. A key distinction is that, while a bacterium always produces two offspring per division, bacteriophage burst sizes are much larger, can themselves be stochastic [12, 37] and, crucially, may strongly depend on lysis time [19, 23]. Here, we elucidate the importance of stochasticity in lysis time and burst size in lytic bacteriophage replication by using agent-based simulations that explicitly model lysis time and burst size distributions and their potential inter-dependencies in two common laboratory competition experiments: serial passage of well-mixed populations and plaque expansion across a bacterial lawn.

We find that a phage’s analytic growth rate in isolation is a poor predictor of the outcome of competition experiments and fails to capture two key features associated with lysis time variability. First, both the serial passage and plaque expansion protocols reward variability in lysis time more than expected: a “noisier” phage is able to outcompete a faster growing, less noisy phage. Second, when lysis times are tightly distributed, we observe an effect we name “population resonance”, in which one phage displays a significant fitness advantage over a competitor with a much greater growth rate in isolation. Both of these phenomena arise from the resource-limited nature of the phage environment and reveal distinct strategies that a phage can adopt to outlive its viral competitors.

## 2 Methods

### 2.1 Lytic bacteriophages can be described by seven life history parameters

The lytic cycle (Fig. 1a) begins with a phage particle (virion) adsorbing to the surface of a susceptible cell. The virion then injects its genetic material (DNA or RNA, depending on the phage species), which hijacks the cell’s molecular machinery. The cell is forced to synthesise many copies of the phage’s proteins and genome, before finally producing lysis proteins, called holins and lysins, which cause the cell to lyse (burst), releasing the assembled virions [48, 49]. Virions then seek new prey through passive diffusion, and may stochastically denature.

**Figure 1:**
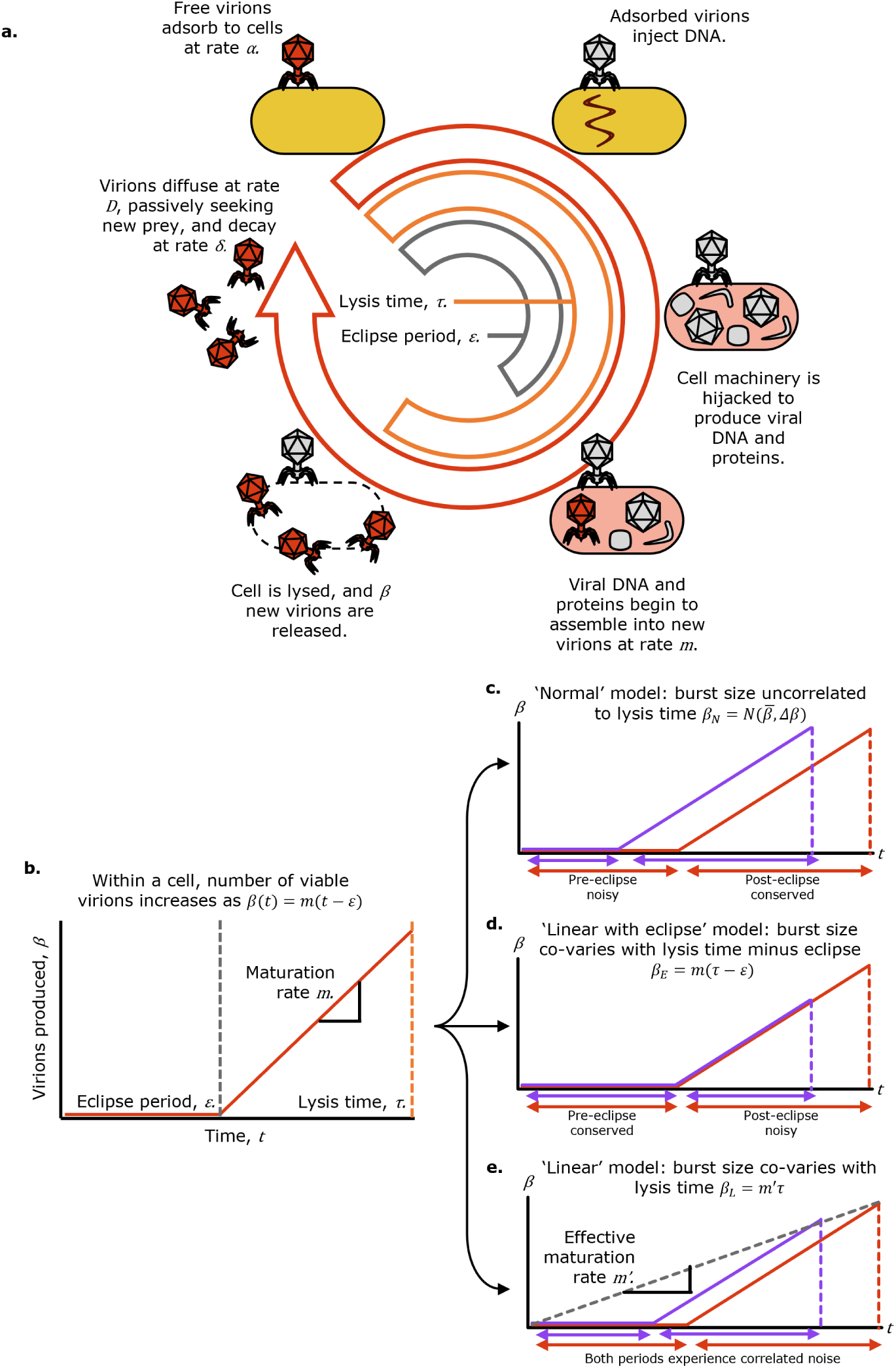
**a.** Schematic of bacteriophage lytic cycle, characterised by seven life history parameters. **b.** Assembly curve, showing the number of viable virions within an infected cell as a function of time: initially zero until the eclipse period elapses, then increasing linearly. **c** to **e.** Assembly curves, each showing the effect of a faster-than-average lysis cycle (purple) versus the average (red), assuming the cycle is faster because the eclipse period is shorter (**c**), the post-eclipse is shorter (**d**), or both are proportionally shorter (**e**).

To compare and predict the action of different bacteriophage strains, we employ a widely-used model [23, 25, 27, 28, 33, 50] in which a bacteriophage is described by seven life history parameters (table 1 and Fig. 1a).

**Table 1:** Viral life history parameters, their units and definitions.

|  |  |
| --- | --- |
| $\alpha$ | Adsorption rate [$\text{ml min}^{-1}$]: rate at which free virions bind to susceptible cells. The chance for a virion to find a bacterium and adsorb to it in time $dt$ is $\alpha B dt$ for bacterial concentration $B$. |
| $\beta$ | Burst size [-]: number of new virions produced in each lysis event. |
| $\tau$ | Lysis time [min], also called ‘latent period’: time between a virion first adsorbing to a healthy cell and cell lysis. |
| $\delta$ | Decay rate [$\text{min}^{-1}$]: rate at which free virions become unviable. |
| $D$ | Diffusion rate [$\mu\text{m}^2\text{s}^{-1}$]: rate at which free virions diffuse through the medium. |
| $\varepsilon$ | Eclipse period [min]: time between a virion first adsorbing to a healthy cell, and the first viable new virion being assembled. |
| $m$ | Maturation rate [$\text{min}^{-1}$]: rate at which virions are assembled inside the cell, following the eclipse period. |

Importantly, these life history parameters do not vary independently, as physics and biology impose both interdependencies and absolute limits. For instance, to release more virions at lysis (greater burst size, *β*), a cell must undertake virion production for longer (greater lysis time, *τ*) or assemble virions faster (greater maturation rate, *m*). We model this dependency through equation 1. This equation has previously been used to describe the assembly of T7 virions by prematurely and externally lysing host cells [19], as well as to relate the lysis times and burst sizes of a number of mutants derived from the same ancestral strain of bacteriophage *λ* [23]. In line with this body of work, we define the relationship between a bacteriophage’s burst size *β* and lysis time *τ* as:

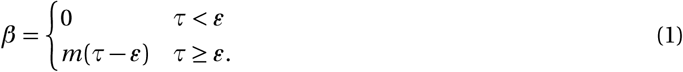

According to this model, the number of viable virions within a cell increases linearly with time, after some initial eclipse period (Fig. 1b). This assumption is consistent with experimental evidence [19, 37] showing that viral components are produced almost linearly until lysis occurs, due to saturation of the host cell’s ribosomes. If lysis times were very long, we expect that this linear relationship might break down, with the assembly curve likely approaching a sigmoid, as the cell’s total available amino acids and nucleotides become the limiting factor (yield-limitation), rather than its ribosomes (rate-limitation). Since no experimental data is currently available in these regimes, here we will focus on the rate-limited case, though all the methods used here could be extended to a logistic model of virion assembly.

While in principle all three parameters relating burst size and lysis time, *m*, *τ* and *ε*, could evolve, it is typically assumed [23] that *m* and *ε* are set by some absolute biological constraints (for a specific bacterial genotype in specific environmental conditions): the production rate *m* is as fast as biologically possible and the eclipse time *ε* is as short as biologically possible. In other words, a larger value of *m* and smaller value of *ε* would always lead to a fitness advantage, so, if the phage could evolve to further increase *m* or decrease *ε*, it would already have done so. The possibility of cell-to-cell variation in *m* is discussed in section 2.2.

As a result, the only remaining parameter that can be subject to selection is *τ*: equation 1 introduces a strategic decision between producing fewer offspring quickly or more offspring slowly. The consequences of these strategies on a bacteriophage’s population-level growth rate, as well as how variability in lysis time implies variability in burst size are discussed in the following section.

### 2.2 Variation in lysis time implies correlated variation in burst size

Assuming virions are produced according to equation 1, there are several possibilities as to how stochasticity in lysis time (*τ*) may create correlated variation in burst size (*β*).

i. If lysis time stochasticity arises purely from variation in the duration of the eclipse period and no variability in production time is observed, no correlated variation in burst size is expected (Fig. 1c). Burst size may still vary from cell-to-cell owing to intrinsic stochasticity in maturation rate, but, crucially, burst size and lysis time will be uncorrelated at the single-cell-single-virion level. We label this possibility the ‘normal’ model. In particular, since in this model lysis time and burst size are uncorrelated, it also represents a null-model where no trade-off between lysis time and burst size is present.
ii. If lysis time stochasticity arises purely from variation in the post-eclipse period of active virion production, then given the lysis time of an individual virion, we can predict the burst size of that virion by applying equation 1 directly (Fig. 1d). We label this the ‘linear with eclipse’ model, or just ‘eclipse’ model.
iii. If lysis time stochasticity affects both the eclipse and post-eclipse proportionately, the relative variation in burst size will be equivalent to the relative variation in lysis time (e.g., 20% reduction in lysis time will lead to 20% reduction in burst size, Fig. 1e). We label this the ‘Linear’ model.

In the following, we consider each of these three models as a biological possibility, and determine in which cases variability in lysis time is advantageous. We focus on variability in lysis time specifically, as variability in burst size alone has little to no effect on a bacteriophage’s ability to propagate itself or clear a cell culture (Fig. S1).

### 2.3 Bacteriophage growth rate in isolation can be predicted by numerical integration

To provide a theoretical expectation for the effect of lysis time variability for the three different model discussed above, we consider a bacteriophage with lysis time distribution *p* (*τ*), and burst size *β* (*τ*), and predict its growth rate under the assumption that bacteria are abundant, and therefore: (a) adsorption is instantaneous, and (b) the probability of adsorbing to an already-infected cell is vanishingly small. Under these assumptions, the population of bacteriophages as a function of time (*V_t_*) is

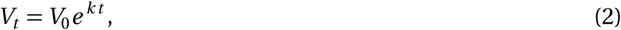

where the effective growth rate *k* is given by

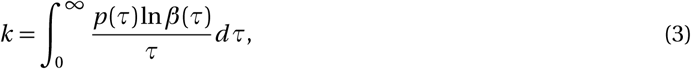

which can be computed numerically for any distribution *p* (*τ*). The extension of this result to a low-density bacterial culture in which adsorption is not fast relative to lysis time can be found in the SI and Fig S11.

### 2.4 Agent-based simulation of two-phage competition experiments

The numerical calculation of the effective growth rate *k* derived above provides a measure of phage selective advantage under the assumption that resources (bacteria) are so abundant that phages do not meaningfully compete against each other. In real case scenarios, including laboratory experiments, phages always interact, if nothing else because of competition for resources. To assess the competitive fitness advantage of a phage population under different stochastic models, we implement two types of agent-based simulations, mimicking two distinct common laboratory protocols used in the study of bacteriophage evolution: serial passage (spatially unstructured)[14, 51] and plaque expansion across a bacterial lawn (spatially structured)[52, 53]. The former is characterized by a total well-mixed volume *Vol* of 10*^−^*^5^ ml (carrying capacity 100,000 cells) initialized with 10,000 uninfected cells at random points in their cell cycle. The latter is modelled as a one-dimensional series of spatial ‘demes’, each corresponding to a length *𝛥x* of 10*µ*m, and volume *Vol* of 10*^−^*^7^ ml (carrying capacity 100 cells), initialized with 100 uninfected cells at random points in their cell cycle. Note that we model a bacterial lawn, not a biofilm and as such all cells are considered to be metabolically active. While the volumes and cell counts used in both simulations are much lower than would be used in a typical laboratory experiment, our model and results are independent of absolute scale (Fig S6), and the values have been chosen to reproduce experimentally observed adsorption curves in well-mixed cultures [8] and plaque expansion speeds on bacterial lawns [25, 54]. We can therefore think of these simulations as representing a subsection of a larger experiment.

Each bacterial agent is characterized by two internal clocks, one representing its own cycle of growth and division and the other representing a count down to lysis, as well as an infection state: ‘healthy’, ‘infected (wild type)’ or ‘infected (mutant)’, and an initially empty list of adsorbed virions. Phage populations are modelled as counters *V_i_* that monitor how many free phages of a specific type are present at any point in time and space. In each simulation time-step, *𝛥t*, ‘bacterial growth’, ‘adsorption’, ‘infection’, ‘lysis’, ‘decay’ and ‘diffusion’ substeps occur (Fig. 2).

**Figure 2:**
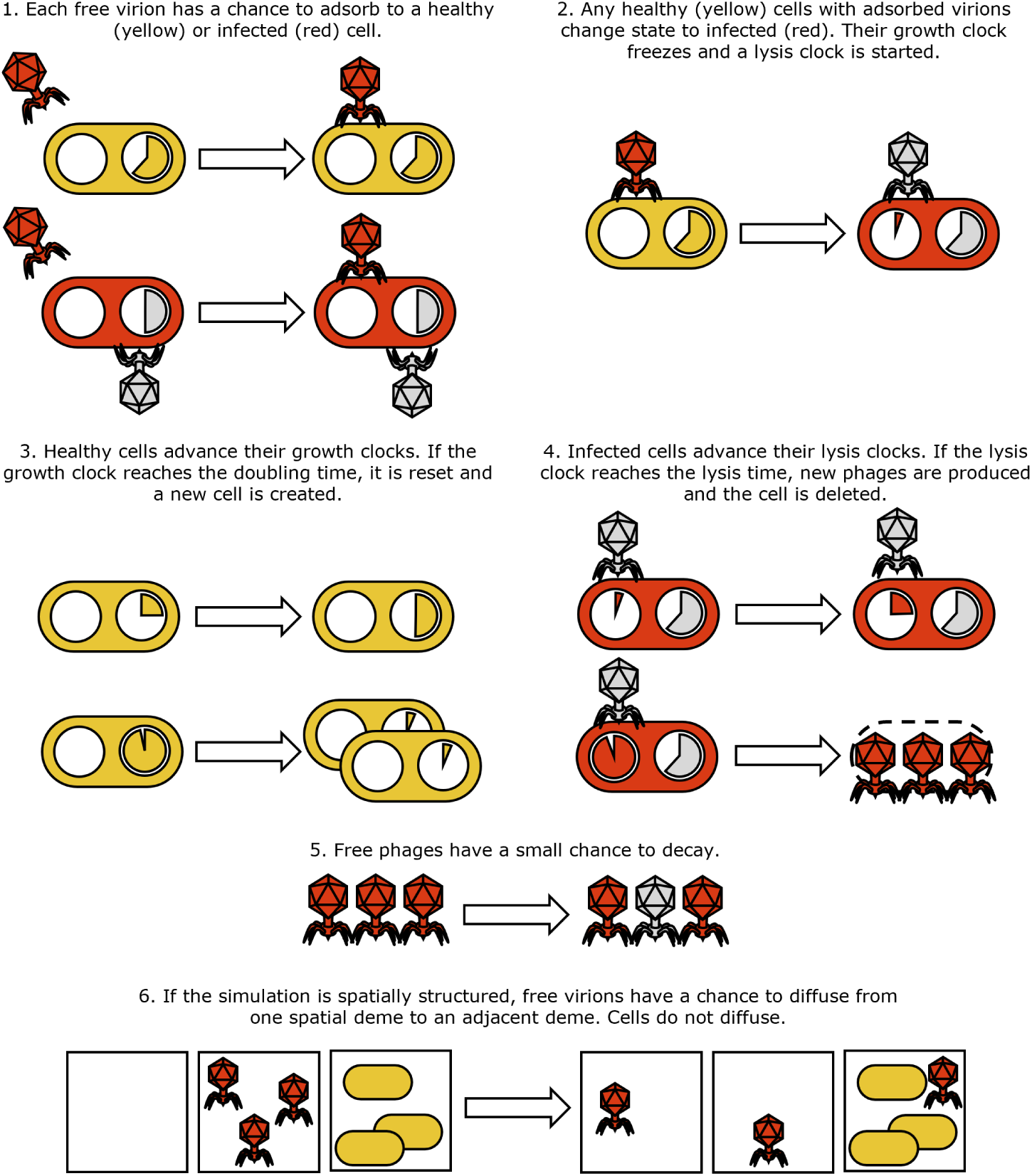
Schematic description of the agent-based model’s implementation. Healthy cells and progressing growth clocks are pictured yellow. Infected cells, viable virions and progressing lysis clocks are red. Virions that have injected their DNA, or that have decayed are pictured grey, as well as bacterial growth clocks which have halted due to infection.

**Adsorption** (Fig. 2 step 1): the number of adsorption events for each phage strain is drawn from a Poisson distribution with mean *α*(*B* + *I*)*V_i_ 𝛥t/Vol*, where *B* is the local number of uninfected bacteria, *I* is the total local number of infected bacteria, and *Vol* is the local volume. A bacterial target, sampled randomly and with replacement, is then associated to each of these events. Adsorptions have a chance to fail proportional to the number of phages already adsorbed to the target cell, representing the finite number of receptor sites each cell possesses. The phage population counter is then updated depending on the number of successful adsorption events.

**Infection** (Fig. 2 step 2): Uninfected bacteria that have at least one adsorbed phage move to an ‘infected (wild type)’ state or ‘infected (mutant)’ state according to the nature of the adsorbed phages. In the case where a cell has adsorbed phages of both types, the type of infected state is chosen binomially using the proportion of adsorbed phages of each type. Once a cell becomes infected, the lysis time is drawn from a normal distribution *У* (*τ*, *𝛥τ*) where *τ* and *𝛥τ* correspond to the lysis time mean and std of the infecting phage, the lysis-clock starts, and the growth-clock is paused. We note that in this setup, there is no benefit to a bacteriophage that adsorbs to an already-infected cell at a later time step, since the first successful infection immediately and permanently determines the cell’s fate. We use the term “super-adsorption” to refer to a virion binding to an already-infected cell, as the secondary virion is removed from the pool but gains no benefit, as the status of the cell is unchanged. This contrasts the phenomenon of “super-infection” [30], in which the secondary virion is able to secure a share of cellular resources, and thus a share of the viral progeny.

**Growth** (Fig. 2 step 3): Uninfected bacteria advance their growth-clocks. If the growth clock completes its cycle, the cell replicates. A new growth cycle length is drawn for both mother and daughter cells from a normal distribution.

**Lysis** (Fig. 2 step 4): infected cells advance their lysis-clocks. If the lysis clock completes its cycle, the cell is deleted, and a number of free phage of the appropriate type are added to their corresponding population counter. Depending on the stochasticity model, this number may be drawn from a distribution, or calculated from the total lysis time.

**Decay** (Fig. 2 step 5): each phage strain’s population counter is reduced by a number drawn from a Poisson distribution with mean *δV 𝛥t*.

**Diffusion** (only for plaque expansion, Fig. 2 step 6): the number of migrating phages from one deme to its neighbours is drawn from a Poisson distribution with mean *DV 𝛥t/*(2*𝛥x* ^2^) and the corresponding phage population counters are appropriately updated.

**Treadmill** (only for plaque expansion, Fig. 6a): to allow the infection front to propagate for longer without having to simulate a very large number of demes, we take a ‘sliding window’ approach. If the deme with the largest number of free virions (total across both pools) is more than 30 demes from the leftmost deme (300*µ*m, or 50% of the total spatial extent *X* of the simulation), the leftmost deme is deleted, all other demes are shunted one position to the left, and a new deme is created at the right hand edge of the simulation.

**Transfer** (only for serial passage, Fig. 4a): if the total number of bacteria falls below 100 (1% of its initial value), we perform a 1000X dilution, so that every remaining bacterium (infected or uninfected) and free phage has 1 in 1000 chance of being transferred to a fresh volume containing 10,000 new uninfected cells. The dilution factor of 1000 [14] and decision to transfer after lysis of the host culture (rather than at fixed time intervals) [51] are both consistent with contemporary evolution experiments.

Both protocols are initialised with 2 pools of 100 phage each, one labelled ‘wild type’, the other labelled ‘mutant’. In the ‘plaque expansion’ all phages are initially placed in the leftmost deme. The serial passage simulations run either until one phage pool outnumbers the other 70:30, at which point it is declared the winner, or for a maximum of 600 minutes. The plaque expansion simulations always run for 180 minutes. The parameter values used in both these simulations are as follows (tables 2 to 5).

**Table 2:** Parameters used in serial passage simulations.

| Serial passage parameters |  |  |
| --- | --- | --- |
| $Vol$ | $10^{-5}$ | Volume [ml] |
| $T$ | 600 | Maximum simulated time [min] |
| $dt$ | 0.01 | Timestep [min] |
| $B_0$ | 10,000 | Initial bacteria count [cells] |
| $C$ | 100,000 | Carrying capacity [cells] |
| $V_0$ | 100 | Initial bacteriophage count [virions per strain] |

**Table 3:** Parameters used in plaque expansion simulations. *The deme volume is important only in determining the rate of adsorption of bacteriophages - on average *αB*/*Vol* phages absorb per unit time.

| Plaque expansion parameters |  |  |
| --- | --- | --- |
| $Vol$ | $10^{-7}$ | Effective volume* [ml per deme] |
| $T$ | 180 | Total simulated time [min] |
| $dt$ | 0.01 | Timestep [min] |
| $X$ | 600 | Total simulated length [μm] |
| $dx$ | 10 | Deme length [μm] |
| $B_0$ | 100 | Initial bacteria count [cells per deme] |
| $C$ | 100 | Carrying capacity [cells per deme] |
| $V_0$ | 100 | Initial bacteriophage count [virions per strain] |

**Table 4:** Bacteria parameters used in all simulations. Values were chosen to agree with published values for *E. coli*, under ideal conditions, but do not represent any particular strain [55]. \**A*_max_ was set to 100, aside from those experiments in which super-adsorption was disabled by setting it to 1.

| Bacteria parameters |  |  |
| --- | --- | --- |
| $\bar{\tau}_D$ | 20 | Mean doubling time [min] |
| $\Delta\tau_D$ | 2 | Doubling time std [min] |
| $A_{\max}$ | 100 or 1* | Maximum number of adsorbed virions [virions] |

**Table 5:** Bacteriophage parameters used in numerical analysis and simulations. Values were chosen to agree with a range of published values for bacteriophage T7, but do not represent any particular strain [8, 36, 44, 51, 56]. *This parameter was either varied within a range, or assigned the value given here. **These Values were chosen such that, in all 3 models, a mean lysis time *τ* of 17 minutes corresponds to a burst size *β* of 150. ^†^*δ* was set to 0.01, aside from those experiments in which decay was disabled by setting it to 0.

| Bacteriophage parameters |  |  |
| --- | --- | --- |
| $\alpha$ | $3 \times 10^{-9}$ | Adsorption rate [$\text{ml min}^{-1}$] |
| $p(\tau)$ | $\mathcal{N}(\bar{\tau}, \Delta\tau)$ | Lysis time probability distribution, taken to be a normal distribution (or delta function in the limit $\Delta\tau \rightarrow 0$) |
| $\bar{\tau}$ | 17 or varied* | Mean lysis time [min] |
| $\Delta\tau$ | 2.5 or varied* | Lysis time std [min] |
| $\varepsilon$ | 9.5** | Eclipse period [min], used in ‘linear with eclipse’ burst size model. |
| $m$ | 20** | Maturation rate [virions $\text{min}^{-1}$], used in ‘linear with eclipse’ burst size model. |
| $m'$ | 8.82** | Effective maturation rate [virions $\text{min}^{-1}$], used in ‘linear’ burst size model. |
| $\bar{\beta}$ | 150** | Mean burst size [virions], used in ‘normal’ burst size model. |
| $\Delta\beta$ | 50 | Mean burst size std [virions], used in ‘normal’ burst size model. |
| $\delta$ | 0.01<sup>†</sup> | Decay rate [$\text{min}^{-1}$] |
| $D$ | 4 | Diffusion rate [$\mu\text{m}^2\text{s}^{-1}$] |

## 3 Results

### 3.1 Numerical calculations provide theoretical expectations for optimal lysis times

In section 2.3, we describe how a bacteriophage’s effective growth rate *k* may be numerically predicted. In Fig. 3a, we report this prediction as a function of mean lysis time for a standard deviation of either 0 or 2.5 minutes, respectively, for the three burst size models. Parameters were chosen to be similar to those of bacteriophage T7, and can be found in table 5. An analysis of the sensitivity of our results to these choices can be found in Figs S12 and S13. In the following: (i) the eclipse period *ε* was taken to be 9.5 minutes, (ii) under all three models, a lysis time *τ* of 17 minutes leads, by construction, to a burst size *β* of 150, and (iii) the lysis time distribution, *p* (*τ*), used is Gaussian, and therefore symmetric. Asymmetric lysis time distributions are considered in Figs S2 to S5.

**Figure 3:**
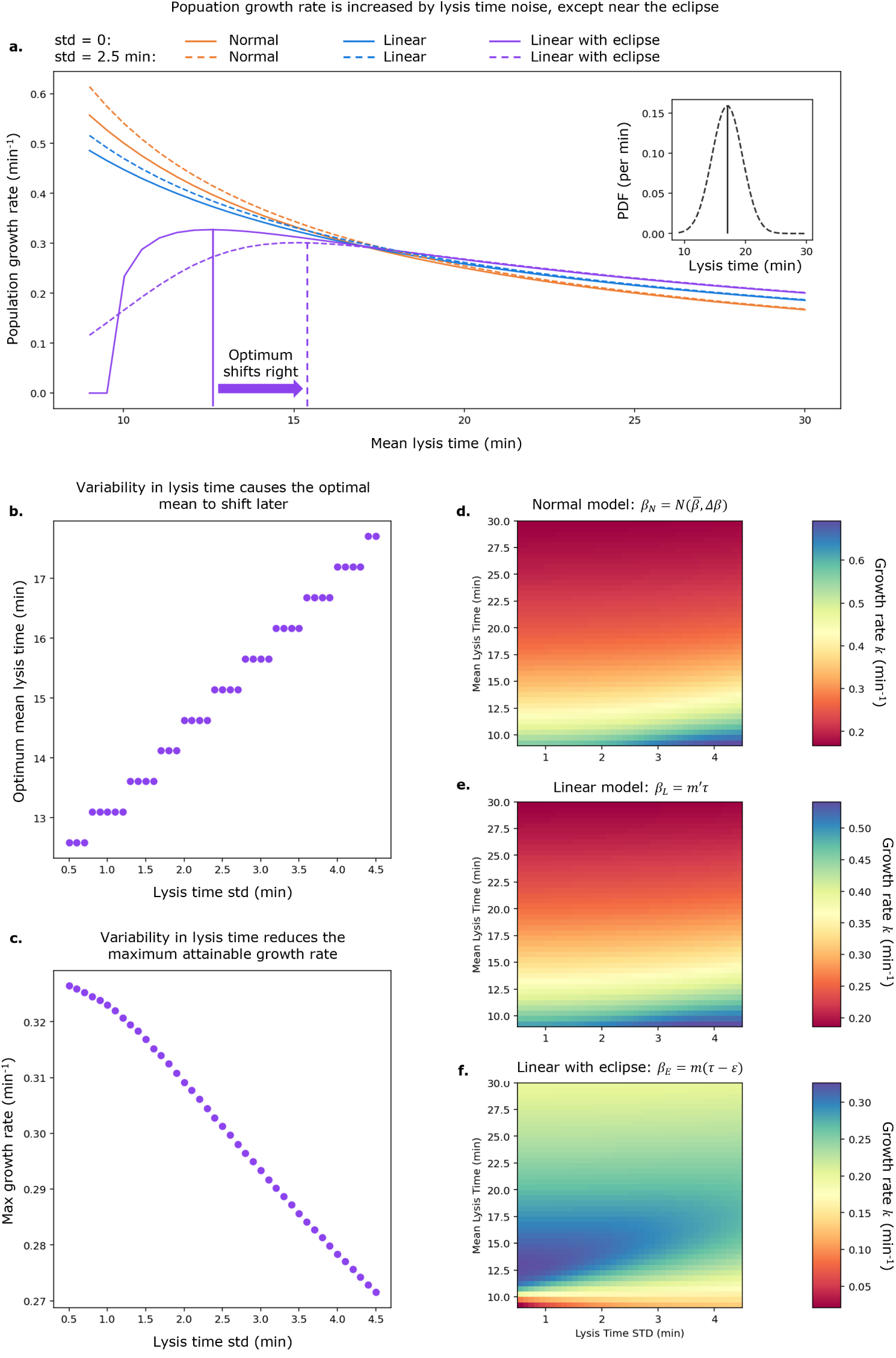
Dependence of bacteriophage growth rate on lysis time under optimal conditions.**a.** Bacteriophage effective growth rate *k* as a function of mean lysis time, with either 0 minutes or 2.5 minutes standard deviation in lysis time. Inset: example lysis time distributions with mean 17 minutes, std 0 (solid line) or 2.5 minutes (dashed line). **b.** Optimal lysis time under ‘linear with eclipse’ model as a function of lysis time std. **c.** Effective growth rate achieved at optimal lysis time, under ‘linear with eclipse’ model, as a function of lysis time std. **d.** to **f.** Effective growth rate as a function of both mean lysis time and lysis time std for ‘normal’, ‘’linear‘, and ‘linear with eclipse’ models respectively.

For the deterministic case (std = 0 min), the effective growth rate *k* decreases with increasing lysis time under the ‘normal’ and ‘linear’ models. By contrast, the ‘linear with eclipse’ model exhibits an optimum lysis time value due to the trade-off between long lysis times leading to very slow replication and short lysis time being unproductive because of the eclipse period. This preference for intermediate lysis times has been observed experimentally in serial passage experiments [14]. We emphasise that the fact that shorter lysis times lead to proportionately smaller bursts, as in the ‘linear’ model, is not sufficient to create an optimal lysis time under exponential growth conditions: the smaller burst size is always more than compensated for by the faster lysis cycle. The existence of this optimum depends entirely on a non-zero eclipse period *ϵ*.

When variability in lysis time is introduced, both the ‘normal’ and ‘linear’ models display an increase in growth rate compared to their deterministic counterparts. Mathematically, this feature can be explained by the convexity of the growth rate function: the mean of a symmetric sample about a given point will yield a greater value than the point itself. Importantly, the variation is symmetric in lysis time. Symmetric variation in lysis rate would cause a decrease in growth rate [57], since the function would no longer be convex. Conversely, the ‘linear with eclipse’ model, which features a concave function for short lysis times, is characterized by a reduction in growth rate with the optimal value shifting rightward (Fig. 3 b,c). Under this model, the noisier a phage’s lysis cycle is, the longer its lysis time should be and the lower the phage maximum growth rate is. This phenomenon holds across a large range of values, as shown in Fig. 3 d,e,f.

Many other probability distributions can be used to investigate the role of noise. A biologically intuitive choice for lysis time distribution is the positively-skewed Erlang distribution, as it describes the stochasticity in waiting time from a series of *n* (shape) consecutive reactions, each occurring with rate *λ* (rate). An Erlang with shape *n* = 1 is simply an exponential distribution. We find that the behaviour of the effective growth rate is qualitatively similar to that obtained using the Gaussian distribution (Fig. S2): ‘Normal’ and ‘Linear’ models favour short mean lysis time and large variance (low shape), while the ‘Linear with Eclipse’ model favours an intermediate optimal mean lysis time and small variance (high shape). A similar trend is also observed for the Weibull distribution (Fig. S3). To look specifically at the effect of asymmetry in the distribution, we have also considered a skew normal distribution (Figs. S4, S5). Interestingly, we see that, for given mean lysis time, negative skew is beneficial in the ‘Normal’ and ‘linear’ models, as it provides additional weight to short lysis times, while positive skew is beneficial in the ‘eclipse’ model, as its short left tail limits the occurrence of very short unproductive lytic cycles. Given the qualitative similarities between the different distributions, in the following, we focus on the easily interpretable Gaussian distribution.

### 3.2 Serial passage competition experiments reveal winning strategies beyond growth rate maximisation

In section 2.4, we described a stochastic, agent-based model of the bacteriophage lytic cycle, which allows us to explicitly model the consequences of variability in a bacteriophage’s burst size and lysis time. We start by using this model to investigate how the phage fitness landscape is affected by (i) the different types of trade-off between lysis time and burst size encoded in our models (Fig. 1c-e), and (ii) the level of noise in lysis time. We consider two bacteriophage populations, labelled “wild type” and “mutant”, and assign them both the same burst size model (‘normal’, ‘linear’, or ‘linear with eclipse’) and the same standard deviation in lysis time (0.01 or 2.5 minutes). We then vary the mean lysis time of each between 10 and 30 minutes in one-minute increments, and compete the two phage populations against each other through a serial passage assay (Fig.4 a, section 2.4). (Fig. 4b, initial MOI 0.01 per phage species, 0.02 total).

**Figure 4:**
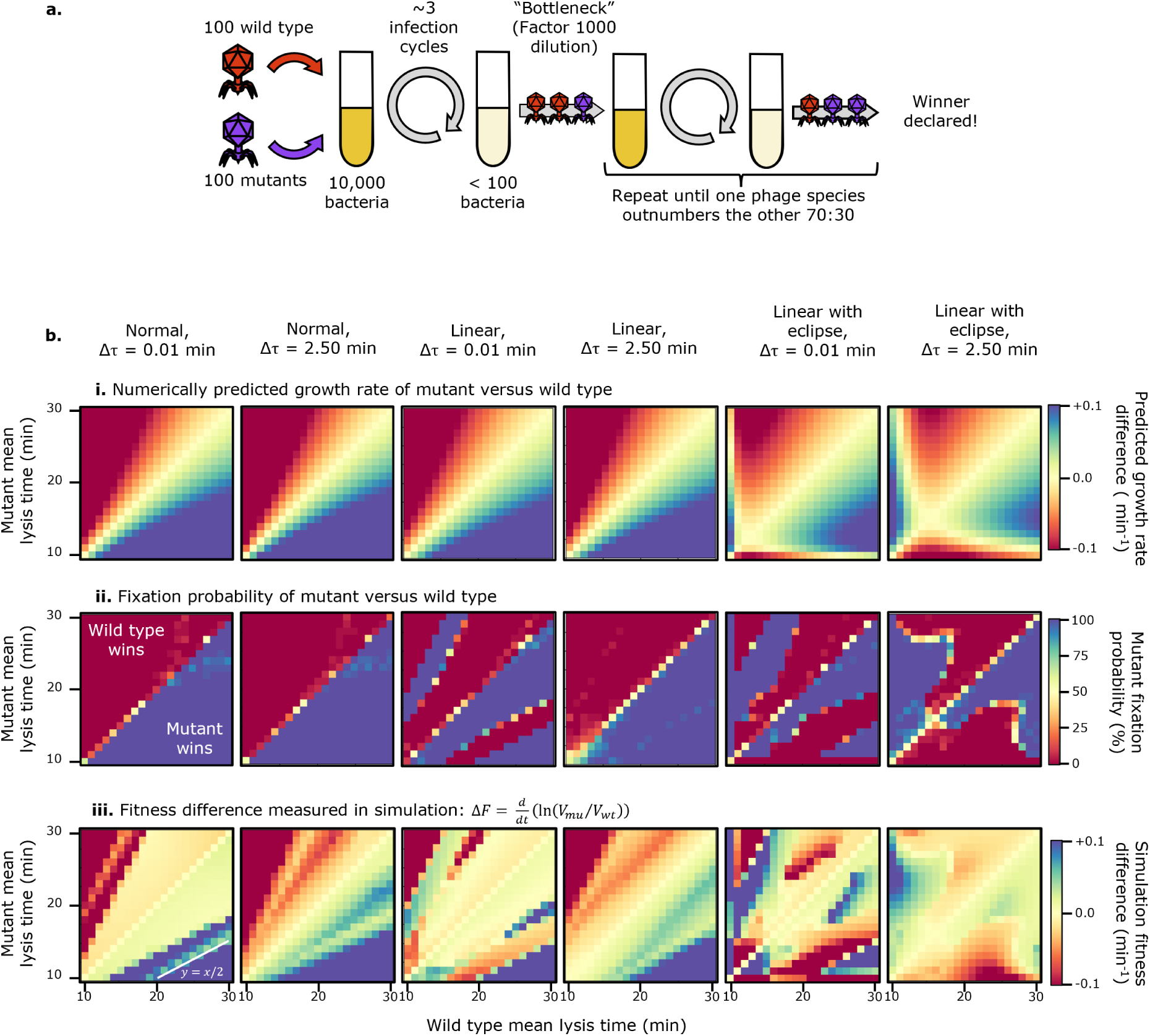
Results of serial passage competition experiment, competing a wild type against a mutant phage with different mean lysis times. **i.** Difference in numerically predicted growth rate, using equation 3. **ii.** Mutant fixation probability in stochastic simulation. **iii.** Fitness difference in stochastic simulation. Note that in the first and third rows, the visual scale has been set to saturate at *±*0.1.

For each set of simulation parameters, we quantify competitive fitness using three alternative measures: (i) the numerically predicted difference in effective growth rate *k* (defined in eq. 3) between the two phages (Fig. 4b i); (ii) the mutant fixation probability defined as the fraction of twenty independent runs in which the mutant fixes in the population (see section 2.4, Fig. 4 b ii); (iii) the rate of fixation of the mutant (positive values) or wild-type (negative values) strain (Fig. 4b iii), defined as

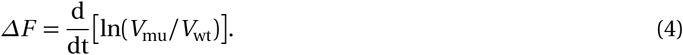

*𝛥F* simplifies to the numerical difference in growth rates *k* in the limiting case where resources are abundant and the two phage populations are growing exponentially (derivation in SI section **??**).

In every case red corresponds to the wild type displaying a fitness advantage and blue to the mutant displaying a fitness advantage. By construction, the plots are antisymmetric about *y* = *x*, therefore in the discussion below we will focus on the bottom-right half of the diagram without loss of generality.

Starting with the leftmost column, corresponding to the ‘normal’ model with narrow lysis time distribution (0.01 minutes standard deviation), the top row (numerically predicted growth rate) replicates the results in Fig. 3 showing that short lysis times lead to a fitness advantage. The middle row (mutant fixation probability) corroborates this numerical result. The bottom row, (mutant fixation rate), reveals an interesting finding. First, we see that while in the majority of parameter space the rate of fixation agrees in sign to the relative growth rate (top row), the rate of fixation increases much more slowly as we move away from the *y* = *x* line. This result reflects how the competition for resources (unaccounted for in the relative growth rate) dramatically screens the fitness advantage that a fast lysing phage would have when grown in isolation (Fig. S10, S14). A similar result was also observed in ref [30] and highlights how phage fitness estimates based on growth in isolation can be misleading and overestimate the fitness (dis)advantage of a population.

The second unexpected result is the appearance of a “stripe” corresponding to intermediate values (green) of fitness advantage within an otherwise highly advantageous region of parameter space (blue). Along this stripe, the mutant lyses cells faster than the wild type and reaches fixation as we would expect. However, it reaches fixation more slowly (in more serial passage cycles) than in the surrounding parameter space.

Close investigation of these cases reveals that they arise due to the limited number of cells available (Fig. S8a). The faster-growing mutant experiences a substantial “dead time” after completing its final round of lysis, just before the transfer step takes place. During this period, the only cells remaining are those infected by the (slower-lysing) wild type. The mutant virions, therefore, have no way to further proliferate, and are only subject to decay or unproductive super-adsorption (“super-adsorption” rather than “super-infection”, as the secondary virion is removed from the pool but gains no benefit, as the status of the cell is unchanged). As a result, by the time the wild type finishes its final round of lysis, there are fewer mutant virions left to survive the dilution.

In line with this explanation, the lower edge of the stripe, highlighted with a white line, corresponds to *y* = *x/*2: the longest possible dead time (and thus the least advantageous parameters for the mutant phage in this region) would occur when the end of the mutant’s final round of lysis happens just after the start of the wild type’s final round. When the mutant’s lysis time becomes half of that of the wild type, it will be able to begin a new round of lysis at the same time as the wild type’s final round, with minimal dead time (Fig S8b).

As additional support to our explanation, we repeat all simulations in Fig. 4b but under “conservative” conditions, in which free virions do not decay and super-adsorption cannot occur (Fig. S10). As expected, under these conditions the stripe disappears, as the mutant phage suffers no penalty from spending “dead time” outside the cell. We refer to this phenomenon as “population resonance”, since it occurs only for specific ratios *τ*_mu_*/τ*_wt_ whose values depend on the number of lytic cycles between dilutions, which are themselves set by the initial multiplicity of infection (MOI) (Fig. S14).

The second column (‘normal’ model, 2.5 mins lysis time std) qualitatively agrees with the first. The only quantitative difference is that the stripe caused by population resonance appears blurred. Since here lysis times for both phages are more variable, lysing events are less synchronised, and the resonance effect is weaker.

The third and fourth columns of Fig.4b correspond to the ‘linear’ model (*β* = *m^𝐹^τ*). When compared to the normal model, the principle difference is that the population resonance effect is stronger. The fixation probability plots (second row) show that, in the 0.01 min std case, population resonance is strong enough to flip which phage reaches fixation. In the 2.5 min case this is not true, as de-synchronisation in lysis cycles weakens the resonance effect, but slight deviations from 100% fixation are visible as a pale blue line in the otherwise uniform block of blue.

The reason why population resonance is stronger under the ‘linear’ model than under the ‘normal’ model is that in the ‘linear’ model longer lysis times confer larger bursts. Hence, while the faster-growing mutant is losing virions during the dead period, the slower-growing wild type benefits from a larger final burst. This longer lysis cycle and larger burst size would not be a winning strategy under exponential growth conditions, as shown in the theoretical expectation for the effective growth rate (top row). However, in the resource-limited conditions of a serial passage, forcing an opponent to lose virions to super-adsorption and decay while taking time to produce more offspring can be a winning strategy. This is in effect a weaponisation of a long lysis time.

Finally, the two rightmost columns on figure 4b correspond to the ‘linear with eclipse’ model (*β* = *m* (*τ−ε*)). The topography of the fitness landscape is altered by the introduction of a critical point corresponding to the optimum lysis time, visible in the top row as the intersection of two (yellow) contours of zero fitness difference and corresponding to the maxima identified in Fig. 3: at lower lysis time values in the low std case, and shifted to larger values in the higher std case. At the same time, the population resonance effect becomes even stronger than under the ‘linear’ model, as longer lysis times imply even greater benefits in burst size.

The fixation probability and the fixation rate plots for the 2.5 min std case (middle and bottom rows, rightmost column) shows that the population resonance stripes are now impactful enough to remain present even with greater lysis time variation, and hence less synchronised lysis cycles, but are “rounded off” by the noise in lysis time. Indeed, a very similar shape can be obtained by applying a Gaussian blur to the corresponding plots in the 0.01 min std case.

In summary, our results reveal the presence of population resonance as a novel effect that arises when two bacteriophage populations in competitions are characterized by specific relative lysis times. It is strongest when lysis time is tightly distributed, and stronger the more the trade-off between lysis time and burst size penalizes short lysis times. According to our simulations, population resonance allows one bacteriophage to outcompete another phage that would grow faster in isolation. One factor which makes population resonance easier to observe is our decision to carry out a transfer once 99% of the initial bacteria have been lysed (as in [51]), as opposed to after some fixed time interval (as in [19]). If a fixed interval were used, and this interval were longer than the time taken to lyse all cells in the culture, the only difference would be an extended period of virion decay between the lysis of the last cell and the transfer step. If both the wild type and mutant have the same decay rate (as in our case), the relative proportion of wild-type and mutant phage would not change, and the resonance is expected to hold in the same position. If, however, the transfer were carried out before the majority of cells had been lysed, additional resonance effects would occur, corresponding to fitness discontinuities as a phage’s lysis time aligns with specific fractions of the dilution interval, producing a more complex fitness landscape (Fig S9).

### 3.3 Serial passage experiments suggest lysis time noise is beneficial even at the cost of reduced growth rate

Having shown that noise in lysis time meaningfully affects which of two competing bacteriophage strains reaches fixation, we now explore whether variability in lysis time can be under selective pressure and quantify its direct effects on competitive fitness. We repeat the serial passage competition experiment described in section 3.2, keeping the wild type phage’s mean lysis time and std constant (17 min and 2.5 min, respectively), while varying those of the mutant (Fig. 5).

**Figure 5:**
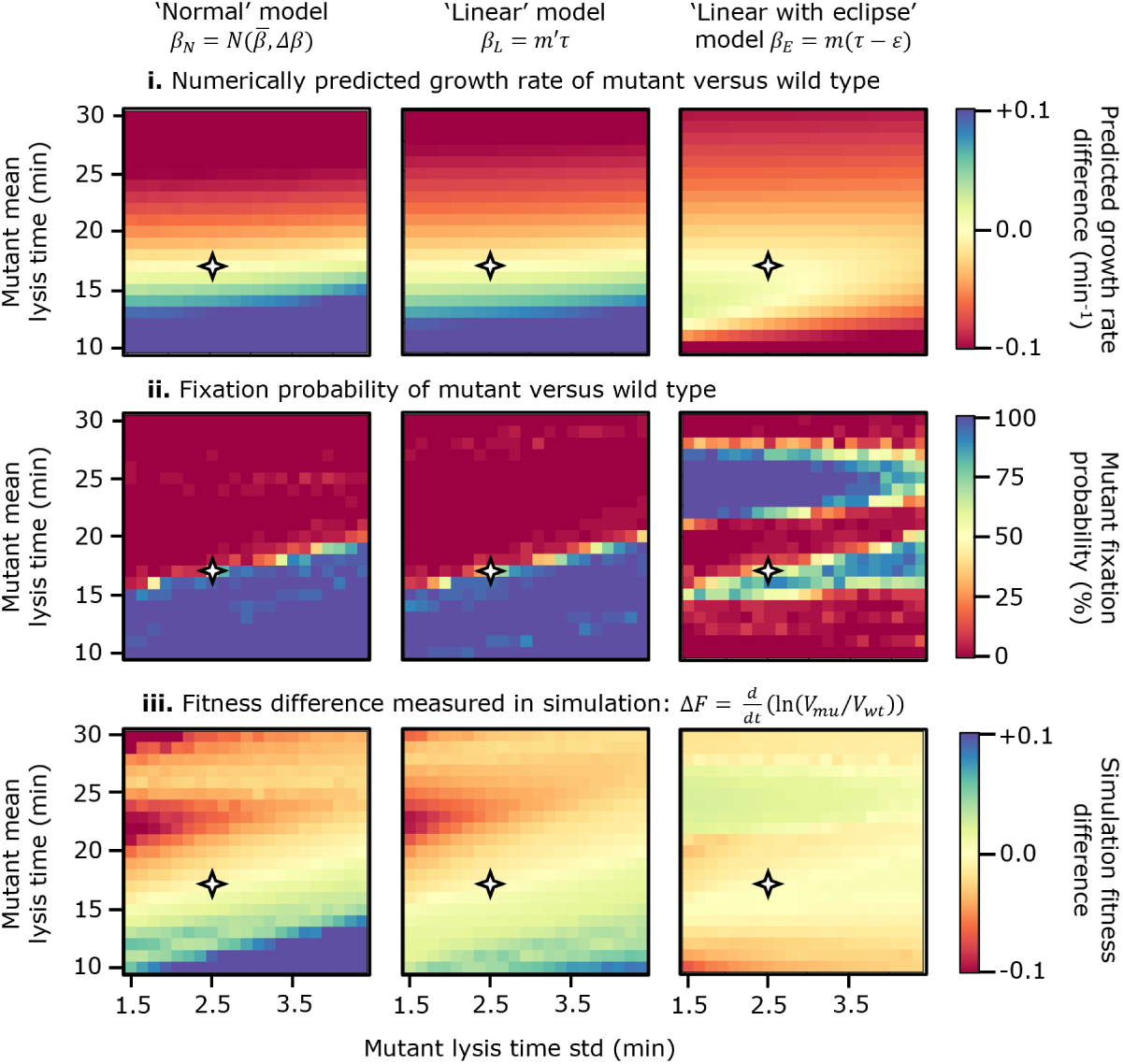
Results of serial passage competition experiment, in which a wild type phage with lysis time mean 17 min, std 2.5 min competes against a mutant with varied lysis time mean (y-axis) and std (x-axis). **i.** Difference in numerically predicted growth rate, using equation 3. **ii.** Mutant fixation probability in stochastic simulation. **iii.** Fitness difference in stochastic simulation. Note that in the first and third rows, the visual scale has been set to saturate at *±*0.1. Axes are scaled such that each pixel step in the y direction corresponds to a 1 minute change in mean lysis time, while each pixel step in the x direction corresponds to an equivalent fractional change in lysis time std versus the wild type value: 2.5/17 minutes. The four-point star on all panels correspond to the point in parameter space at which both phages have exactly the same parameters.

The first two columns, corresponding to the ‘normal’ and ‘linear’ models, are qualitatively similar to each other. In both cases, the theoretical growth rate difference (top row) suggests that a mutant with a shorter lysis time should grow faster than the wild type. The yellow contour of equal fitness has a slightly positive gradient, meaning that a phage should see an equal gain in fitness by either reducing its mean lysis time by 1 minute, or increasing its lysis time std by approximately 3 minutes.

The middle row (fixation probability) agrees with the overall claim that shorter lysis times are advantageous, but the slope of the equal-fitness contour is greater than in the top row, suggesting that, under the resource-limited conditions of the serial passage, each 1 minute decrease in lysis time mean is competitively equivalent to an increase of approximately 1 min in lysis time std. Since this increase in gradient similarly holds for both the ‘normal’ and the ‘linear’ models, it cannot be understood in terms of the trade-off between lysis time and burst size characteristic of the latter, but it is the manifestation of an intrinsic benefit of stochasticity under resource-limited conditions.

An explanation for this result is the following. In exponential growth, one bacteriophage strain will outcompete the other if its growth rate is consistently higher, therefore its lysis cycles must be fast on average. In the serial passage experiment, bacteria are limited and vastly outnumbered by the phage towards the end of each passage (MOI approaching 100). Here, a bacteriophage strain does not need to have short lysis cycles on average, rather it needs enough short lysis cycles to secure a larger share of the available bacteria, which can be achieved with large variation around the mean.

The rate of fixation (bottom row) provides a finer quantification of these results, as we see that the mutant fitness disadvantage at high mean lysis times predicted by growth in isolation can be overcome by increasing lysis time std. For a given std, the fitness difference as a function of mean lysis time displays a non-monotic behaviour. This is a result of the population resonance effect described in the previous section. Importantly, while increasing noise decreases the relative advantage emerging from the resonance effect, it also broadens the region of parameter space in which it is relevant (Fig. S15i).

The rightmost column of Fig. 5 corresponds to the ‘linear with eclipse‘ model. Here, the numerical growth rate prediction (top row) points to a small optimal region of parameter space (short lysis time, low std, lime green) below and to the left of the equivalence point, in which the mutant should grow faster than the wild type. This reflects the finding in section 2.3, and the fact that the wild type phage is already close to the optimum. Interestingly, the mutant fixation probability plot (second row) shows a different picture, in which two distinct regions of parameter space result in mutant fixation. The region towards the top of the plot is the result of population resonance (when super-adsorption and decay are disabled this region disappears, Fig. S16).

The high-fitness region at the bottom of the plot, which in the other models extends to the *x* -axis, is here instead bounded from below by the presence of the eclipse period: if the mean lysis time of the mutant is too close to the eclipse period, the mutant will not be competitive. This result agrees with numerical prediction (top row). The fitness difference in simulations, however, disagrees with numerical predictions in that the majority of this high fitness region is to the right of the equivalence point (larger stds), rather than below and to the left. This is because, as mentioned previously, when resources are sufficiently limited that only a small number of lysis cycles are possible, the optimal strategy is not to maximise steady-state growth rate, but rather to carry out fast early lysis cycles, which secure a majority share of the available resources, followed by slower but more productive later lysis cycles. An intermediate mean lysis time and larger lysis time std make this possible. The lysis time std cannot be made too great, or the phage risks lysing before or shortly after the eclipse period, producing nothing. Hence, the lower edge of this high-fitness region is slightly upward sloped, favoring longer mean lysis times when the std is large.

In summary, we find that serial passage rewards lysis time variability more than numerical growth rate predictions would suggest (Fig S15), including some cases in which numerical growth rate wrongly predicts that variability would be detrimental. We reason that this is due to fast lysis cycles being disproportionately advantageous under resource-limited conditions as they ensure a greater share of the total, limited, pool of resources (bacteria). In the ‘linear’ and ‘linear with eclipse’ models, larger lysis time std allows for both shorter lysis cycles that produce virions quickly to claim the available resources, and longer, more productive later cycles that generate more virions. It should be stressed that this winning strategy is achieved only probabilistically: an individual phage is equally likely to cause a faster-than-average first lysis cycle as to cause a slower-than-average cycle. If the phage could sense the culture MOI and ‘decide’ to lyse rapidly when MOI is low and slowly when MOI is high, such as through a lysis-inhibition mechanism [58], this would be an even more competitive strategy.

### 3.4 Plaque expansion assay does not exhibit population resonance and results in weaker selective pressure on lysis time

We now move to a second standard phage growth assay, a plaque expansion across a bacterial lawn, which we simulate as a one dimensional series of well-mixed demes (Fig. 6), as detailed in the Methods section. Free virions, but not bacteria, are able to diffuse between adjacent demes (Fig. 6a). The initial MOI (per deme) is 1 per phage species, 2 total, in the leftmost deme, and 0 elsewhere, representing the typical local densities of viral particles in an initial inoculation of a clearing.

**Figure 6:**
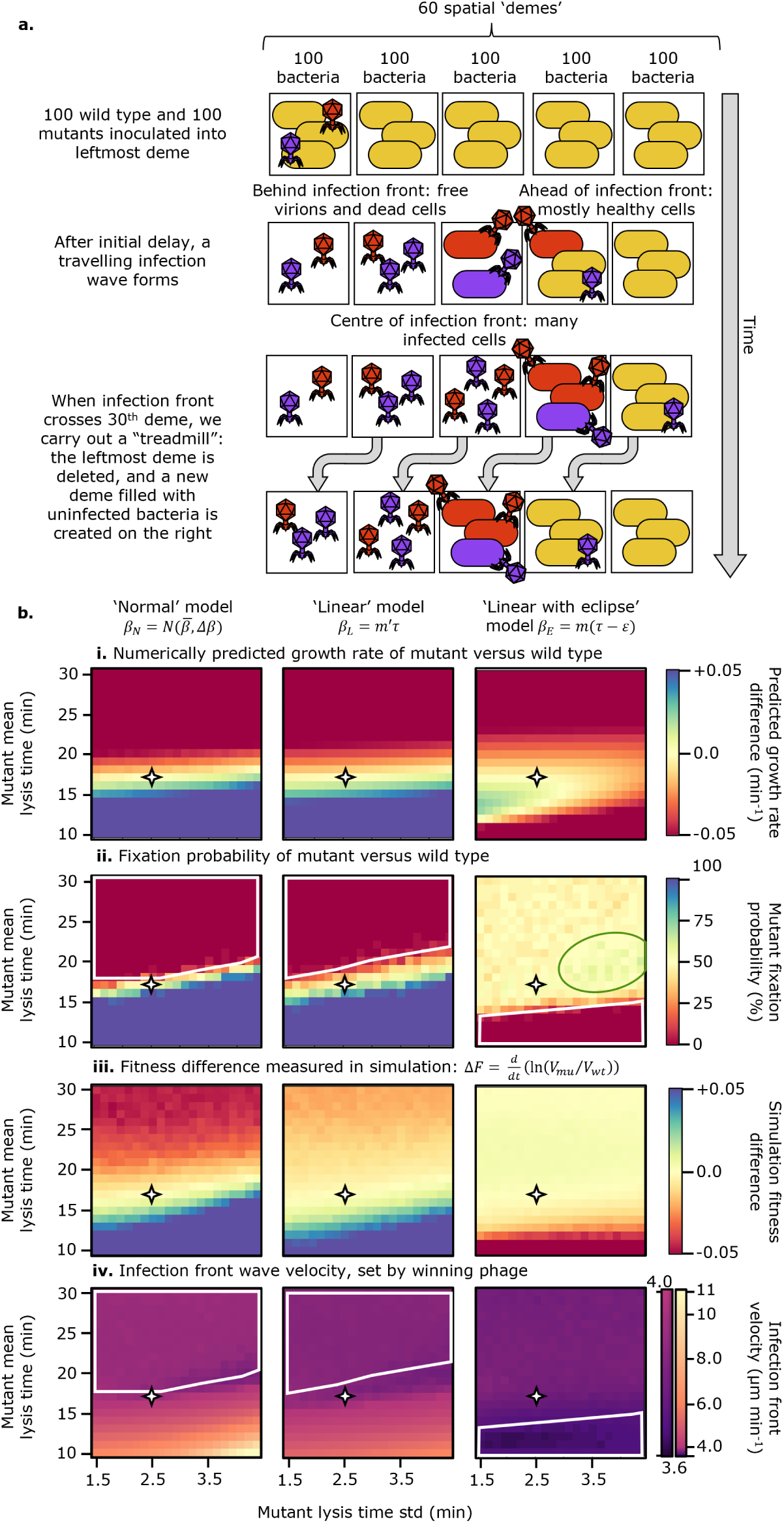
Plaque expansion simulation. **a.** Plaque expansion experiment schematic. **b.** Results of plaque expansion competition experiment, in which a wild type phage with lysis time mean 17 min, std 2.5 min competes against a mutant with varied lysis time mean (y-axis) and std (x-axis). **i.** Difference in numerically predicted growth rate, using equation 3. **ii.** Mutant fixation probability in stochastic simulation. **iii.** Fitness difference in stochastic simulation. **iv.** Infection front velocity. Note that in the first and third rows, the visual scale has been set to saturate at *±*0.05. Axes are scaled such that each pixel step in the y direction corresponds to a 1 minute change in mean lysis time, while each pixel step in the x direction corresponds to an equivalent fractional change in lysis time std versus the wild type value: 2.5/17 minutes. The four-point star on all panels corresponds to the point in parameter space at which both phages have exactly the same parameters. The white polygons in rows ii and iv outline the parameter space in which the wild type reaches fixation.

The data in the top row is identical to that in Fig. 5, with a different colour range to make it directly comparable with the data from the simulations of plaque expansions. For the ‘normal’ and ‘linear’ models, the results share some common features with the serial passage case. Particularly, the line of equal fitness has a steeper gradient in the stochastic simulation plot (second row) than the predicted growth rate plot (top row), suggesting that lysis time variation is still rewarded more strongly than growth rate alone would predict. This is not surprising, as the typical MOI in plaque expansion is very high, making resources even more limited.

The fitness difference plots (third row) are markedly different to those produced by serial passage. First, the population resonance effect is no longer present, since virions no longer experience a “dead period” waiting for new bacteria to be added and lysis cycles quickly become desynchronised because of diffusion. Secondly, fitness (dis)advantages are overall smaller in magnitude here than in the serial passage case. In particular, a mutant with a lysis time mean larger than the wild type’s is able to persist for much longer and, vice-versa, a fast lysing mutant will take more time to fix.

The final row shows the infection wave’s velocity, which is determined by the winning phage. A mutant which produces a much faster infection front will quickly overtake and outcompete the wild type phage. The white polygon corresponds to the region of parameter space in which the wild type phage reaches fixation, and therefore we expect the wave velocity to be constant. Mutants with shorter mean lysis times and greater lysis time std produce faster infection waves, but infection waves are significantly slower in the linear model (max approx. 6 *µ*m/min) than in the normal model (max approx. 11 *µ*m/min), as a result of the trade-off between lysis time and burst size.

The third column corresponds to the ‘linear with eclipse’ model. As in the serial passage experiment, the optimal region of parameter space in which the mutant is able to reach fixation (second row, green ellipse) is not located where the numerical growth rate would suggest (top row). Its shift towards high std values strongly supports the idea that lysis time noise is beneficial in resource-limited environments.

The third row (fitness difference) and fourth row (infection wave velocity) show a general loss of contrast across the majority of the parameter space. This suggests that when a strong trade-off between lysis time and burst size is present (second and third columns), the plaque expansion puts little (second column, ‘linear’ model) to almost no (third column, ‘linear with eclipse’ model) selective pressure on lysis time, regardless of its distribution, as any benefit coming from lysing faster is compensated by the lower burst size.

In summary, a spatially structured environment removes any population resonance effect, and masks the selective pressure on lysis time compared to a serial passage setup. In line with this, we also find that, while noise in lysis time still confers a detectable fitness advantage, its overall impact is less significant, especially when a lysis time-burst size trade-off is present.

## 4 Discussion

### 4.1 Variability in lysis time as an evolvable trait with significant impact on fitness

Clonal bacteria growing under identical conditions display significant heterogeneity in gene expression due to intrinsic biochemical noise [29]. It is therefore expected that bacteriophages, which heavily rely on their host’s molecular machinery, are subject to similar unavoidable stochasticity in their replication. The extent and pervasiveness of such noise in a phage’s life history parameters cannot be measured by traditional phenotypic assays based on bulk population measurements, and as such, they have remained inaccessible until recently [12, 36, 37]. As a result, the vast majority of experimental and theoretical work aimed at quantifying phage ecology and evolution has focused on fitness effects linked to average values. Recent advances in single-cell measurements of phage-bacteria interactions [12, 37] and more careful analyses of population assays [11] are finally enabling an experimental quantification of variability in life history parameters, inspiring parallel theoretical work.

Far from “noise” being simply an effect to which bacteriophages are subject, the degree of variability in a phage’s life history parameters has recently been shown to be, at least partially, under the phage genetic control [35, 36, 45], raising questions as to when and why such noise might be favoured by natural selection, or, alternatively, selected against. This parallels a broader trend in evolutionary biology, which sees stochasticity as not just influencing evolution, but, in some cases, driving it [59, 60].

In this work, using extensive agent-based simulations, we have investigated how phenotypic stochasticity in lysis time and burst size can confer fitness (dis)advantages to a phage population in two commonly used experimental assays: serial passage [14] and plaque expansion [61]. Overall, we observe two surprising phenomena. First, variability in lysis time is generally an advantageous trait, even when it reduces growth rate in isolation. Second, when lysis cycles remain tightly synchronised, a novel effect we name “population resonance” may occur, in which a slower growing phage effectively weaponises a longer lysis cycle to create a sink for competing phages.

### 4.2 Lysis time variability is more advantageous than expected and its effects depend on the lysis time-burst size trade-off

One of the fundamental differences between noise in division time in bacteria and noise in lysis time in phages is the trade-off between the length of the latent period *τ* and the burst size *β* (unlike for bacteria, which will always replicate into two daughter cells upon division). For bacteria, it is known that variability in doubling time is advantageous, as long as it is symmetric [46]. The same is true for single-celled eukaryotes such as budding yeast [62]. By contrast, our results show that, for phages, the details of the trade-off between lysis time and burst size have strong influence on the fitness landscape and on how noise modulates it. Surprisingly, a linear correlation between burst size and lysis time results in a fitness map that is very similar to the one generated in the absence of any trade-off in both serial passage and plaque expansion scenarios. In both situations, we find a strong selection towards fast lysis cycles, whether this occurs by decreasing the mean lysis time, or by increasing the variability in lysis time so that more infection cycles are completed rapidly. For a non-zero optimal lysis time to emerge, a non-productive eclipse period has to be introduced in the trade-off. In this case, theory predicts that an optimal lysis time with minimal variation will produce the fastest growth rate. Interestingly, we find that, in both the serial passage and a plaque expansion competitions, variability in lysis time remains advantageous, even where it reduces population growth rate.

We find that the primary selector for such phenotypic variability is resource limitation. Most laboratory settings, and likely wild environments, are characterized by periods of time or regions of space with very high MOI, where many phage virions compete for few host cells. In these conditions, noise in lysis time appears to confer increased resilience to a phage population. By producing virions throughout time and not in synchronized large bursts, variable lysis times can establish a more steady flow of virions, so that susceptible cells can be infected whenever and wherever they become available. This effect becomes crucial when free virions are subject to rapid loss, e.g., because of super-adsorption. Importantly, the desynchronisation achieved by lysis time variability is not equivalent to that achieved by a lower adsorption rate: a low adsorption rate inevitably reduces growth rate and competitive advantage because it increases the time spent by the phage searching for a host (see predicted growth rates in Fig. 4 vs Fig. S11). By contrast, a variable lysis time with high adsorption rate allows any released phage to immediately infect a host a soon as it becomes available. Reduced adsorption rate may still be beneficial if prolonged periods of very high MOI are common, since it reduces a bacteriophage’s chance to be wasted through super-adsorption, but it would come at the cost of slowing down infection when susceptible cells do become available.

A loose analogy can be drawn between variability in lysis time and the stochastic emergence of bacterial ‘persisters’, the fraction of bacterial cells which enter a state of near-zero growth, even when resources are abundant [63]. Both strategies reduce the population’s growth rate while conditions remain ideal, but make it more resilient to massive loss under stressful conditions: extended periods of high MOI, or the application of antibiotics respectively.

### 4.3 Population resonance increases the number of viable lytic strategies

Our second key observation is the novel phenomenon we name ‘population resonance’, in which a phage which grows slowly in isolation is able to outcompete a faster-growing phage in serial passage experiments. In order for population resonance to occur, three conditions are necessary. First, lysis cycles must be comparatively synchronised. This means variation in lysis time must be small, and that adsorption must occur quickly, which implies that the cell culture must be relatively dense. Second, a trade-off between lysis time and burst size must be present, so that longer lysis times lead to larger burst sizes. Third, free virions must be subject to some decay, for instance via super-adsorption to previously infected cells. Since resources (susceptible bacteria) are limited, each phage will undergo a finite number of lysis cycles before the cell population is exhausted. If the slower-lysing phage’s lysis time value is just less than a multiple of the faster-lysing phage’s lysis time, for example 19 and 10 minutes respectively, it will be able to adsorb to the last uninfected cells present, forcing the final burst of fast-lysing phage to undergo a long “dead period” in which the free virions are subject to decay. By the time the slower-lysing phage completes its final lysis cycle, its offspring will outnumber what is left of the fast-lysing phages.

Combining our understanding of population resonance with the competitive advantage offered by lysis time noise provides an interpretation for the complex fitness landscape created by the serial passage experiment, particularly in the case of the ‘linear with eclipse’ model. We find that this assay favours two opposite strategies: “patient” and “rapacious”. The “patient” strategy uses a relatively slow mean lysis time with little variability, so that the phage can exploit the population resonance effect and effectively “outlive” the competing faster-lysing phage. The “rapacious” strategy favours shorter lysis time with high variability, so that the phage quickly removes potential host cells for the slower-lysing competing phage and relies on the occasional longer cycles for productive infections. Both the “rapacious” and the “patient” strategies are able to outcompete a “prudent” strategy that hinges on maximising growth rate in isolation (intermediate mean lysis time, very small std) because of the competition for limited resources (cells) between two phage populations. The efficacy of a similarly rapacious strategy was observed in [64], in which a fast-migrating, fast-lysing phage was able to out-compete a slower-lysing prudent phage, which grew more quickly in isolation. Conversely, lysis-inhibition [58] and lysogeny [65, 66] which are both extant strategies in wild type phages, may be thought of as extensions of the “patient” strategy: when MOI is high, bursts are delayed as released virions would be likely lost to unproductive-super adsorption.

It would be interesting to test experimentally which strategy would actually preferentially emerge in an evolutionary experiment, since a serial passage is a well established laboratory protocol. While the outcome will be sensitive to the availability of mutations and potential epistasis, we expect that the rapacious strategy (short lysis, large variability) will typically be more robust, since the presence and location of population resonance, which is leveraged by the patient strategy, strongly depends on the details of the competing phage, the number of cells at the beginning of each serial passage, and the initial MOI (see Figs. S10, S14, S16).

Turning to the plaque expansion assay, unexpectedly, we find that introducing an eclipse period in the lysis time-burst size trade-off almost completely flattens the fitness landscape, as long as lysis time is larger than the eclipse period. This result implies that there should be almost no selective pressure on lysis time or its variability in an evolutionary experiment performed on a bacterial lawn, possibly explaining why several studies performing such experiments predominantly find mutations related to adsorption [64, 67]. Interestingly, our finding also suggests an indirect way to identify the presence of a strong trade-off between burst size and lysis time. A plaque assay competition dominated by fast-lysing phages would indicate the absence of a strong trade-off. In contrast, a plaque competition dominated by slow, highly productive phages could indicate the presence of a strong trade-off between lysis time and burst size, and, likely, the presence of a strongly conserved eclipse period. It is important to highlight that the plaque expansion assay is distinct from the infection of a bacterial biofilm. In plaque expansion, phages propagate across a metabolically homogenous bacterial lawn. By contrast, in a bacterial biofilm we would expect significant heterogeneity in the metabolic states of the bacteria [68, 69]. Extending the model presented here to a bacterial biofilm would need to account for heterogeneity in nutrient availability, and therefore metabolic rate, as well as the presence of distinct bacterial phenotypes [68, 69].

### 4.4 Population resonance has implications in both laboratory and wild environments

Population resonance, leveraged by the “patient” viral strategy, depends on synchronised infection events. In the serial passage protocol, this is achieved by the synchronous initial inoculation of phages into bacterial culture, followed by the synchronous inoculation into a fresh bacterial culture. While this synchrony is undoubtedly artificially amplified in some laboratory setting, population resonance may have broader applicability to other natural or artificial systems which are periodically flushed and in which bacteria and phages coexist, such as tidal aquatic environments [70], or the or human urinary [71] and gastrointestinal tract [72, 73], if not as the primary driver of selection, then as an important ‘second-order’ consideration. For example, if one wishes to eradicate a pathogenic bacterial strain from the human GI or urinary tract, a cocktail of “rapacious” virulent phages and “patient” phages, potentially either lysogenic or possessing a lysis-inhibition-mechanism, might lead to better outcome. Reciprocally, if population resonance is predominantly confined to a laboratory setting, our results warn against over-interpretation of the emergence of optimal lysis times in experimental evolution, which might simply be the result of the selection protocol employed.

### 4.5 Summary

Predicting how an organism will fare in competition, based on its behaviour in an ideal environment with abundant resources and no meaningful competition, is challenging [74]. Resource investments ranging from toxin production in bacteria [75, 76] to predation-defences in plants [77, 78] reduce growth rate in monoculture, but become vital once competitors or predators are introduced. This phenomenon applies equally to bacteriophage. While phages lack the toxins and trichomes necessary to directly remove their competitors, we have demonstrated that fast, noisy, or in some cases very slow lysis cycles can be weaponised to allow one phage to outcompete another, at the cost of reducing its growth rate in isolation.

Can variability in lysis time be an adapted trait? When no or minimal trade-off between lysis time and burst size exists, our results show that selection will simply push the phage to lyse as quickly as possible. In this case, noisy phages could potentially emerge only if there is a lower limit to how short a deterministic lysis time can be or if it is evolutionary easier, because of availability of mutations or presence of epistatic effects, to increase variance in lysis time rather than decreasing its mean. However, in the presence of an eclipse period, an optimal lysis time value exists. Once the phage reaches this optimum, we expect that the selective pressure will concentrate on optimizing variability in lysis time. Experimentally testing these hypotheses will not only clarify the evolutionary role of noise in phage ecology and evolution, but also reveal which genetic changes can modulate variability in lysis time. Thus far, identified mutations which contribute to lysis time variability seem to hit the genes responsible for the expression of holin proteins, both in *λ* phage [36] and *φX* 174 [45]. As holins are usually among the final genes expressed in the lytic cycle, partially responsible for lysis of the host [79], we expect that noise should affect only the post eclipse period, since virion production is already underway by the time holins are transcribed. Single-cell microfluidic experiments interestingly seem to suggest that the period between the beginning of capsid protein synthesis and lysis is under comparatively tight control, and that the period between adsorption and capsid expression is much more variable in wild-type phage T7 [37]. Taken together, these data suggests that variability in the lytic cycle of a bacteriophage is likely a complex phenomenon with no single source. Further experiments and genetic analysis of ‘noisy’ mutants are needed to deepen our understanding of the intricate interplay between an infecting phage and its host.

## 5 Data availability

The stochastic, agent-based simulation “Stochastic 3.3.2” was written in Python 3, based on earlier computational work [30, 37]. Simulations were executed on the CSD3 high performance computing cluster at University of Cambridge. The code used for the full simulation, minimal “fast” simulation, numerical integration and heat-map plotting is available at https://github.com/FuscoLab/lysis_time_noise.

## 6#Acknowledgements

AS and DF were supported by an ERC Starting Grant/UKRI Horizon Europe Guarantee (EP/Y030141/1). MH was supported by a UKRI EPSRC Doctoral Studentship from the Department of Physics, University of Cambridge (2125180). Simulations in this work were performed using resources provided by the Cambridge Service for Data Driven Discovery (CSD3) operated by the University of Cambridge Research Computing Service (www.csd3.cam.ac.uk), provided by Dell EMC and Intel using Tier-2 funding from the Engineering and Physical Sciences Research Council (capital grant EP/T022159/1), and DiRAC funding from the Science and Technology Facilities Council (www.dirac.ac.uk). No funding bodies played any role in the study design, data collection and analysis, decision to publish, or preparation of the manuscript.

## 7 List of Supporting Information

- SI section 1: Single phage killing curve simulation suggests variability in burst size alone has little to no effect on bacteriophage fitness.
  **–** Fig S1: Results of ‘killing curve’ assays.
- SI section 2: Derivation of fitness difference *𝛥F*.
- SI section 3: Asymmetric and symmetric lysis time distributions exhibit the same qualitative behaviour.
  **–** Fig S2: Erlang distribution numerical results.
  **–** Fig S3: Weibull distribution numerical results.
  **–** Fig S4: Skew normal distribution numerical results, varying skew and mean.
  **–** Fig S5: Skew normal distribution numerical results, varying skew and std.
- SI section 4: Mean-mean serial passage controls.
  **–** Fig S6: Serial passage simulation results at three different absolute scales.
  **–** Fig S7: Serial passage simulation results using two possible definitions of ‘fitness’.
  **–** Fig S8: Close investigation of fitness ‘stripes’ created by population resonance.
  **–** Fig S9: Serial passage simulation results comparing transfer-on-clearing protocol with transfer after fixed 25 minute interval.
  **–** Fig S10: Serial passage simulation results in which super-adsorption, virion decay, and separately cell growth are disabled.
  **–** Fig S11: Serial passage simulation results in low-adsorption regime.
- SI section 5: Mean-mean minimal simulation controls.
  **–** Fig S12: Minimal simulation results for cell doubling times much faster than, approximately equal to, or much slower than phage lysis time.
  **–** Fig S13: Minimal simulation results for additional values of maturation rate *m* and eclipse period *𝗌*.
  **–** Fig S14: Minimal simulation results for cell counts varied over several orders of magnitude.
  **–** SI subsection 5.3: Implementation of minimal simulation.
- SI section 6: Assay comparison reveals weaker selective pressure in plaque assay and broadening of resonance stripes at high lysis time std.
  **–** Fig S15: Assay comparison for serial passage experiment.
- SI section 7: The secondary fitness maximum observed under the ‘linear with eclipse’ model in explained by population resonance.
  **–** Fig S16: Serial passage simulation results for cell counts varied over an order of magnitude, and sperately with super-adsorption disabled.

## References

[1] Anne Chevallereau et al. “Interactions between bacterial and phage communities in natural environments”. In: Nature Reviews Microbiology 20.1 (Jan. 2022). Publisher: Nature Publishing Group, pp. 49–62. ISSN: 1740-1534. DOI: 10.1038/s41579-021-00602-y. URL: https://www.nature.com/articles/s41579-021-00602-y (visited on 01/22/2025).

[2] Charles Schmidt. “Phage therapy’s latest makeover”. In: Nature Biotechnology 37.6 (June 1, 2019). Bandiera_abtest: a Cg_type: Nature Research Journals Number: 6 Primary_atype: News Publisher: Nature Publishing Group, pp. 581–586. ISSN: 1546-1696. DOI: 10.1038/s41587-019-0133-z. URL: https://www.nature.com/articles/s41587-019-0133-z (visited on 01/25/2022).

[3] Review on Antimicrobial Resistance. Tackling Drug-Resistant Infections Globally: Final Report and Recommendations. UK Government and Wellcome Trust, 2016, p. 80. URL: https://amr-review.org/sites/default/files/160525_Final%20paper_with%20cover.pdf (visited on 01/30/2022).

[4] William C. Summers. “The strange history of phage therapy”. In: Bacteriophage 2.2 (Apr. 1, 2012). Publisher: Taylor & Francis _eprint: 10.4161/bact.20757, pp. 130–133. ISSN: null. DOI: 10.4161/bact.20757. URL: 10.4161/bact.20757 (visited on 06/30/2025).

[5] Félix d’Herelle. “An Invisible Microbe that is Antagonistic to the Dysentery Bacillus (Sur un microbe invisible antagonsite des bacilles dysenteriques.” In: Comptes rendus Acad. Sciences 165 (1917), pp. 373–375.

[6] Rui Zhu et al. “In Vitro and In Vivo Antibacterial Efficacy of Bacteriophage Combined with Tigecycline Against Carbapenem-Resistant Klebsiella pneumoniae and Characterization of Phage Resistant Mutants”. In: Frontiers in Cellular and Infection Microbiology 15 (June 26, 2025). Publisher: Frontiers. ISSN: 2235-2988. DOI: 10.3389/fcimb.2025.1610625. URL: https://www.frontiersin.org/journals/cellular-and-infection-microbiology/articles/10.3389/fcimb.2025.1610625/full (visited on 06/30/2025).

[7] A. Vieira et al. “Phage therapy to control multidrug-resistant Pseudomonas aeruginosa skin infections: in vitro and ex vivo experiments”. In: European Journal of Clinical Microbiology & Infectious Diseases 31.11 (Nov. 1, 2012), pp. 3241–3249. ISSN: 1435-4373. DOI: 10.1007/s10096-012-1691-x. URL: 10.1007/s10096-012-1691-x (visited on 06/30/2025).

[8] M. Delbrück. “ADSORPTION OF BACTERIOPHAGE UNDER VARIOUS PHYSIOLOGICAL CONDITIONS OF THE HOST”. In: The Journal of General Physiology 23.5 (May 20, 1940), pp. 631–642. ISSN: 0022-1295. URL: https://www.ncbi.nlm.nih.gov/pmc/articles/PMC2237952/ (visited on 04/17/2023).

[9] E. L. Ellis and M. Delbrück. “THE GROWTH OF BACTERIOPHAGE”. In: The Journal of General Physiology 22.3 (Jan. 20, 1939), pp. 365–384. ISSN: 0022-1295. DOI: 10.1085/jgp.22.3.365.

[10] Albert P. Krueger. “THE SORPTION OF BACTERIOPHAGE BY LIVING AND DEAD SUSCEPTIBLE BACTERIA”. In: The Journal of General Physiology 14.4 (Mar. 20, 1931), pp. 493–516. ISSN: 0022-1295. URL: https://www.ncbi.nlm.nih.gov/pmc/articles/PMC2141123/ (visited on 05/27/2023).

[11] Marian Dominguez-Mirazo et al. “Accounting for cellular-level variation in lysis: implications for virus–host dynamics”. In: mBio 0.0 (July 19, 2024). Publisher: American Society for Microbiology, e01376–24. DOI: 10.1128/mbio.01376-24. URL: https://journals.asm.org/doi/10.1128/mbio.01376-24 (visited on 08/01/2024).

[12] Sherin Kannoly et al. “Single-Cell Approach Reveals Intercellular Heterogeneity in Phage-Producing Capacities”. In: Microbiology Spectrum 11.1 (Dec. 21, 2022). Publisher: American Society for Microbiology, e02663–21. DOI: 10.1128/spectrum.02663-21. URL: https://journals.asm.org/doi/full/10.1128/spectrum.02663-21 (visited on 06/30/2025).

[13] Sílvio B. Santos et al. “Population Dynamics of a Salmonella Lytic Phage and Its Host: Implications of the Host Bacterial Growth Rate in Modelling”. In: PLOS ONE 9.7 (July 22, 2014). Publisher: Public Library of Science, e102507. ISSN: 1932-6203. DOI: 10.1371/journal.pone.0102507. URL: https://journals.plos.org/plosone/article?id=10.1371/journal.pone.0102507 (visited on 06/30/2025).

[14] Sherin Kannoly, Abhyudai Singh, and John J. Dennehy. “An Optimal Lysis Time Maximizes Bacterio-phage Fitness in Quasi-Continuous Culture”. In: mBio 13.3 (Apr. 25, 2022). Publisher: American Society for Microbiology, e03593–21. DOI: 10.1128/mbio.03593-21. URL: https://journals.asm.org/doi/full/10.1128/mbio.03593-21 (visited on 06/30/2025).

[15] Shazeeda Koonjan, Carlos Cardoso Palacios, and Anders S. Nilsson. “Population Dynamics of a Two Phages–One Host Infection System Using Escherichia coli Strain ECOR57 and Phages vB_EcoP_SU10 and vB_EcoD_SU57”. In: Pharmaceuticals 15.3 (Mar. 2022). Number: 3 Publisher: Multidisciplinary Digital Publishing Institute, p. 268. ISSN: 1424-8247. DOI: 10.3390/ph15030268. URL: https://www.mdpi.com/1424-8247/15/3/268 (visited on 04/27/2023).

[16] Katja Šivec and Aleš Podgornik. “Determination of bacteriophage growth parameters under cultivating conditions”. In: Applied Microbiology and Biotechnology 104.20 (Oct. 2020), pp. 8949–8960. ISSN: 1432-0614. DOI: 10.1007/s00253-020-10866-8.

[17] Romain Gallet, Yongping Shao, and Ing-Nang Wang. “High adsorption rate is detrimental to bacteriophage fitness in a biofilm-like environment”. In: BMC Evolutionary Biology 9.1 (Oct. 5, 2009), p. 241. ISSN: 1471-2148. DOI: 10.1186/1471-2148-9-241. URL: 10.1186/1471-2148-9-241 (visited on 05/12/2023).

[18] Daniel Campos, Vicenç Méndez, and Sergei Fedotov. “The effects of distributed life cycles on the dynamics of viral infections”. In: Journal of Theoretical Biology 254.2 (Sept. 21, 2008), pp. 430–438. ISSN: 0022-5193. DOI: 10.1016/j.jtbi.2008.05.035. URL: https://www.sciencedirect.com/science/article/pii/S0022519308002932 (visited on 06/19/2023).

[19] Richard H. Heineman and James J. Bull. “Testing optimality with experimental evolution: lysis time in a bacteriophage”. In: Evolution; International Journal of Organic Evolution 61.7 (July 2007), pp. 1695– 1709. ISSN: 0014-3820. DOI: 10.1111/j.1558-5646.2007.00132.x.

[20] Dongwoo Chae. “Phage-host-immune system dynamics in bacteriophage therapy: basic principles and mathematical models”. In: Translational and Clinical Pharmacology 31.4 (Dec. 2023), pp. 167–190. ISSN: 2289-0882. DOI: 10.12793/tcp.2023.31.e17. URL: https://www.ncbi.nlm.nih.gov/pmc/articles/PMC10772058/ (visited on 06/30/2025).

[21] Anders S. Nilsson. “Cocktail, a Computer Program for Modelling Bacteriophage Infection Kinetics”. In: Viruses 14.11 (Nov. 2022). Number: 11 Publisher: Multidisciplinary Digital Publishing Institute, p. 2483. ISSN: 1999-4915. DOI: 10.3390/v14112483. URL: https://www.mdpi.com/1999-4915/14/11/2483 (visited on 06/30/2025).

[22] Quentin J. Leclerc, Jodi A. Lindsay, and Gwenan M. Knight. “Modelling the synergistic effect of bacteriophage and antibiotics on bacteria: Killers and drivers of resistance evolution”. In: PLOS Computational Biology 18.11 (Nov. 30, 2022). Publisher: Public Library of Science, e1010746. ISSN: 1553-7358. DOI: 10.1371/journal.pcbi.1010746. URL: https://journals.plos.org/ploscompbiol/article?id=10.1371/journal.pcbi.1010746 (visited on 06/30/2025).

[23] Ing-Nang Wang. “Lysis Timing and Bacteriophage Fitness”. In: Genetics 172.1 (Jan. 1, 2006), pp. 17–26. ISSN: 1943-2631. DOI: 10.1534/genetics.105.045922. URL: 10.1534/genetics.105.045922 (visited on 05/12/2023).

[24] Joaquim Fort and Vicenç Méndez. “Time-Delayed Spread of Viruses in Growing Plaques”. In: Physical Review Letters 89.17 (Oct. 8, 2002). Publisher: American Physical Society, p. 178101. DOI: 10.1103/PhysRevLett.89.178101. URL: https://link.aps.org/doi/10.1103/PhysRevLett.89.178101 (visited on 07/04/2025).

[25] J Yin and J S McCaskill. “Replication of viruses in a growing plaque: a reaction-diffusion model.” In: Biophysical Journal 61.6 (June 1992), pp. 1540–1549. ISSN: 0006-3495. URL: https://www.ncbi.nlm.nih.gov/pmc/articles/PMC1260448/ (visited on 03/06/2024).

[26] Jacopo Marchi et al. Stable coexistence and transport of lytic phage infections with migrating bacterial hosts. Pages: 2025.04.18.649568 Section: New Results. Apr. 21, 2025. DOI: 10.1101/2025.04.18.649568. URL: https://www.biorxiv.org/content/10.1101/2025.04.18.649568v2 (visited on 05/22/2025).

[27] Melinda Choua et al. “The effect of viral plasticity on the persistence of host-virus systems”. In: Journal of Theoretical Biology 498 (Aug. 7, 2020), p. 110263. ISSN: 0022-5193. DOI: 10.1016/j.jtbi.2020. 110263. URL: https://www.sciencedirect.com/science/article/pii/S0022519320301181 (visited on 10/05/2022).

[28] Gabor Beke, Matej Stano, and Lubos Klucar. “Modelling the interaction between bacteriophages and their bacterial hosts”. In: Mathematical Biosciences 279 (Sept. 1, 2016), pp. 27–32. ISSN: 0025-5564. DOI: 10.1016/j.mbs.2016.06.009. URL: https://www.sciencedirect.com/science/article/ pii/S0025556416300700 (visited on 06/30/2025).

[29] Michael B. Elowitz et al. “Stochastic Gene Expression in a Single Cell”. In: Science 297.5584 (Aug. 16, 2002). Publisher: American Association for the Advancement of Science, pp. 1183–1186. DOI: 10.1126/science.1070919. URL: https://www.science.org/doi/10.1126/science.1070919 (visited on 07/11/2025).

[30] Michael Hunter and Diana Fusco. “Superinfection exclusion: A viral strategy with short-term benefits and long-term drawbacks”. In: PLOS Computational Biology 18.5 (May 10, 2022). Publisher: Public Library of Science, e1010125. ISSN: 1553-7358. DOI: 10.1371/journal.pcbi.1010125. URL: https://journals.plos.org/ploscompbiol/article?id=10.1371/journal.pcbi.1010125 (visited on 05/29/2023).

[31] Andrés Valdez et al. “Biomechanical modeling of spatiotemporal bacteria-phage competition”. In: Communications Physics 8.1 (Apr. 8, 2025). Publisher: Nature Publishing Group, p. 139. ISSN: 2399-3650. DOI: 10.1038/s42005-025-02078-1. URL: https://www.nature.com/articles/s42005-025-02078-1 (visited on 07/04/2025).

[32] Blessing Emerenini et al. “The effect of environmental factors in biofilm and phage interactions in Agent Based Model”. In: *arXiv* (Feb. 22, 2025). URL: 10.48550/arXiv.2502.18507 (visited on 06/30/2025).

[33] Michael Hunter et al. “Virus-Host Interactions Shape Viral Dispersal Giving Rise to Distinct Classes of Traveling Waves in Spatial Expansions”. In: Physical Review X 11.2 (June 29, 2021). Publisher: American Physical Society, p. 021066. DOI: 10.1103/PhysRevX.11.021066. URL: https://link.aps.org/doi/10.1103/PhysRevX.11.021066 (visited on 06/21/2023).

[34] Alison Cameron et al. Keep Your Frenemies Closer: Bacteriophage That Benefit Their Hosts Evolve to be More Temperate. 2022. URL: https://ecoevorxiv.org/repository/object/3802/download/7521/.

[35] Abhyudai Singh and John J. Dennehy. “Stochastic holin expression can account for lysis time variation in the bacteriophage ”. In: Journal of The Royal Society Interface 11.95 (June 6, 2014). Publisher: Royal Society, p. 20140140. DOI: 10.1098/rsif.2014.0140. URL: https://royalsocietypublishing.org/doi/10.1098/rsif.2014.0140 (visited on 07/04/2025).

[36] John J. Dennehy and Ing-Nang Wang. “Factors influencing lysis time stochasticity in bacteriophage lambda”. In: BMC Microbiology 11.1 (Aug. 2, 2011), p. 174. ISSN: 1471-2180. DOI: 10.1186/1471-2180-11-174. URL: 10.1186/1471-2180-11-174 (visited on 10/05/2022).

[37] Charlie Wedd et al. Single-cell imaging of the lytic phage life cycle in bacteria. Pages: 2024.04.11.588870 Section: New Results. Oct. 22, 2024. DOI: 10.1101/2024.04.11.588870. URL: https://www.biorxiv.org/content/10.1101/2024.04.11.588870v3 (visited on 06/30/2025).

[38] Antrea Pavlou et al. “Single-cell data reveal heterogeneity of investment in ribosomes across a bacterial population”. In: Nature Communications 16.1 (Jan. 2, 2025). Publisher: Nature Publishing Group, p. 285. ISSN: 2041-1723. DOI: 10.1038/s41467-024-55394-5. URL: https://www.nature.com/articles/s41467-024-55394-5 (visited on 07/04/2025).

[39] Kimberly M. Davis and Ralph R. Isberg. “Defining heterogeneity within bacterial populations via single cell approaches”. In: BioEssays 38.8 (2016). _eprint: https://onlinelibrary.wiley.com/doi/pdf/10.1002/bies.201500121, pp. 782–790. ISSN: 1521-1878. DOI: 10.1002/bies.201500121. URL: https://onlinelibrary.wiley.com/doi/abs/10.1002/bies.201500121 (visited on 07/14/2025).

[40] Lingchong You, Patrick F. Suthers, and John Yin. “Effects of Escherichia coli Physiology on Growth of Phage T7 In Vivo and In Silico”. In: Journal of Bacteriology 184.7 (Apr. 2002), pp. 1888–1894. ISSN: 0021-9193. DOI: 10.1128/JB.184.7.1888-1894.2002. URL: https://www.ncbi.nlm.nih.gov/pmc/articles/PMC134924/ (visited on 05/05/2023).

[41] Erin L. Attrill et al. “Individual bacteria in structured environments rely on phenotypic resistance to phage”. In: PLOS Biology 19.10 (Oct. 12, 2021). Publisher: Public Library of Science, e3001406. ISSN: 1545-7885. DOI: 10.1371/journal.pbio.3001406. URL: https://journals.plos.org/plosbiology/article?id=10.1371/journal.pbio.3001406 (visited on 05/15/2023).

[42] Maxime Schwartz. “The adsorption of coliphage lambda to its host: Effect of variations in the surface density of receptor and in phage-receptor affinity”. In: Journal of Molecular Biology 103.3 (May 25, 1976), pp. 521–536. ISSN: 0022-2836. DOI: 10.1016/0022-2836(76)90215-1. URL: https://www.sciencedirect.com/science/article/pii/0022283676902151 (visited on 07/14/2025).

[43] Stephen Tobias Abedon. “Bacteriophage Adsorption: Likelihood of Virion Encounter with Bacteria and Other Factors Affecting Rates”. In: Antibiotics 12.4 (Apr. 2023). Number: 4 Publisher: Multidisciplinary Digital Publishing Institute, p. 723. ISSN: 2079-6382. DOI: 10.3390/antibiotics12040723. URL: https://www.mdpi.com/2079-6382/12/4/723 (visited on 07/04/2025).

[44] James J. Bull, Richard H. Heineman, and Claus O. Wilke. “The Phenotype-Fitness Map in Experimental Evolution of Phages”. In: PLOS ONE 6.11 (Nov. 22, 2011). Publisher: Public Library of Science, e27796. ISSN: 1932-6203. DOI: 10.1371/journal.pone.0027796. URL: https://journals.plos.org/plosone/article?id=10.1371/journal.pone.0027796 (visited on 07/04/2025).

[45] Christopher W Baker et al. “Genetically Determined Variation in Lysis Time Variance in the Bacteriophage X174”. In: G3 Genes|Genomes|Genetics 6.4 (Apr. 1, 2016), pp. 939–955. ISSN: 2160-1836. DOI: 10.1534/g3.115.024075. URL: https://doi.org/10.1534/g3.115.024075 (visited on 11/19/2025).

[46] Mikihiro Hashimoto et al. “Noise-driven growth rate gain in clonal cellular populations”. In: Proceedings of the National Academy of Sciences 113.12 (Mar. 22, 2016). Publisher: Proceedings of the National Academy of Sciences, pp. 3251–3256. DOI: 10.1073/pnas.1519412113. URL: https://www.pnas.org/doi/10.1073/pnas.1519412113 (visited on 05/09/2023).

[47] E. O. Powell. “Growth Rate and Generation Time of Bacteria, with Special Reference to Continuous Culture”. In: Microbiology 15.3 (1956). Publisher: Microbiology Society, pp. 492–511. ISSN: 1465-2080. DOI: 10.1099/00221287-15-3-492. URL: https://www.microbiologyresearch.org/content/journal/micro/10.1099/00221287-15-3-492 (visited on 05/11/2023).

[48] Bálint Kiss et al. “Imaging the Infection Cycle of T7 at the Single Virion Level”. In: International Journal of Molecular Sciences 23.19 (Jan. 2022). Number: 19 Publisher: Multidisciplinary Digital Publishing Institute, p. 11252. ISSN: 1422-0067. DOI: 10.3390/ijms231911252. URL: https://www.mdpi.com/1422-0067/23/19/11252 (visited on 01/10/2023).

[49] Gunther Stent. Molecular Biology of Bacterial Viruses. A Series of Books in Biology 17. United States of America: W. H. Freeman and Company, 1963. 474 pp.

[50] Yongping Shao and Ing-Nang Wang. “Bacteriophage Adsorption Rate and Optimal Lysis Time”. In: Genetics 180.1 (Sept. 1, 2008), pp. 471–482. ISSN: 1943-2631. DOI: 10.1534/genetics.108.090100. URL: 10.1534/genetics.108.090100 (visited on 09/15/2023).

[51] Hai Xu et al. “Biological Characterization and Evolution of Bacteriophage T7-holin During the Serial Passage Process”. In: Frontiers in Microbiology 12 (2021), p. 705310. ISSN: 1664-302X. DOI: 10.3389/fmicb.2021.705310.

[52] Stephen T. Abedon and Rachel R. Culler. “Optimizing bacteriophage plaque fecundity”. In: Journal of Theoretical Biology 249.3 (Dec. 7, 2007), pp. 582–592. ISSN: 0022-5193. DOI: 10.1016/j.jtbi.2007.08.006. URL: https://www.sciencedirect.com/science/article/pii/S0022519307003815 (visited on 07/11/2025).

[53] J Yin. “Evolution of bacteriophage T7 in a growing plaque.” In: Journal of Bacteriology 175.5 (Mar. 1993), pp. 1272–1277. ISSN: 0021-9193. DOI: 10.1128/jb.175.5.1272-1277.1993. URL: https://www.ncbi.nlm.nih.gov/pmc/articles/PMC193211/ (visited on 07/11/2025).

[54] Michael Hunter. “The Role of Virus-Host Interactions in the Evolutionary Dynamics of Bacteriophage Populations”. PhD thesis. University of Cambridge, June 2022. 190 pp.

[55] Mike S. Son and Ronald K. Taylor. “Growth and Maintenance of Escherichia coli Laboratory Strains”. In: Current protocols 1.1 (Jan. 2021), e20. ISSN: 2691-1299. DOI: 10.1002/cpz1.20. URL: https://www.ncbi.nlm.nih.gov/pmc/articles/PMC8006063/ (visited on 07/14/2025).

[56] J. J. Bull and I. J. Molineux. “Predicting evolution from genomics: experimental evolution of bacteriophage T7”. In: Heredity 100.5 (May 2008). Publisher: Nature Publishing Group, pp. 453–463. ISSN: 1365-2540. DOI: 10.1038/sj.hdy.6801087. URL: https://www.nature.com/articles/6801087 (visited on 07/14/2025).

[57] Jie Lin and Ariel Amir. “The Effects of Stochasticity at the Single-Cell Level and Cell Size Control on the Population Growth”. In: Cell Systems 5.4 (Oct. 25, 2017), 358–367.e4. ISSN: 2405-4712. DOI: 10.1016/j.cels.2017.08.015. URL: https://www.sciencedirect.com/science/article/pii/S2405471217303873 (visited on 04/23/2024).

[58] Holly Kloos Dressman and John W. Drake. “Lysis and Lysis Inhibition in Bacteriophage T4: rV Mutations Reside in the Holin t Gene”. In: Journal of Bacteriology 181.14 (July 1999), pp. 4391–4396. ISSN: 0021-9193. DOI: 10.1128/jb.181.14.4391-4396.1999. URL: https://www.ncbi.nlm.nih.gov/pmc/articles/PMC93942/ (visited on 07/11/2025).

[59] Magali Richard and Gaël Yvert. “How does evolution tune biological noise?” In: Frontiers in Genetics 5 (Oct. 28, 2014), p. 374. ISSN: 1664-8021. DOI: 10.3389/fgene.2014.00374. URL: https://pmc.ncbi.nlm.nih.gov/articles/PMC4211553/ (visited on 11/20/2025).

[60] Matjaž Perc and Attila Szolnoki. “Noise-guided evolution within cyclical interactions”. In: New Journal of Physics 9.8 (Aug. 2007), p. 267. ISSN: 1367-2630. DOI: 10.1088/1367-2630/9/8/267. URL: 10.1088/1367-2630/9/8/267 (visited on 11/20/2025).

[61] Elhanan Tzipilevich et al. “Bacteria elicit a phage tolerance response subsequent to infection of their neighbors”. In: The EMBO Journal 41.3 (Feb. 1, 2022), e109247. ISSN: 0261-4189. DOI: 10.15252/embj.2021109247. URL: https://pmc.ncbi.nlm.nih.gov/articles/PMC8804946/ (visited on 11/20/2025).

[62] Bram Cerulus et al. “Noise and Epigenetic Inheritance of Single-Cell Division Times Influence Population Fitness”. In: Current Biology 26.9 (May 9, 2016), pp. 1138–1147. ISSN: 0960-9822. DOI: 10.1016/j.cub.2016.03.010. URL: https://www.sciencedirect.com/science/article/pii/S0960982216301865 (visited on 12/05/2025).

[63] Bram Van den Bergh, Maarten Fauvart, and Jan Michiels. “Formation, physiology, ecology, evolution and clinical importance of bacterial persisters”. In: FEMS Microbiology Reviews 41.3 (May 1, 2017), pp. 219–251. ISSN: 0168-6445. DOI: 10.1093/femsre/fux001. URL: 10.1093/femsre/fux001 (visited on 11/25/2025).

[64] Benjamin Kerr et al. “Local migration promotes competitive restraint in a host–pathogen ’tragedy of the commons’”. In: Nature 442.7098 (July 2006). Publisher: Nature Publishing Group, pp. 75–78. ISSN: 1476-4687. DOI: 10.1038/nature04864. URL: https://www.nature.com/articles/nature04864 (visited on 07/09/2025).

[65] Cristina Howard-Varona et al. “Lysogeny in nature: mechanisms, impact and ecology of temperate phages”. In: The ISME Journal 11.7 (July 2017). Publisher: Nature Publishing Group, pp. 1511–1520. ISSN: 1751-7370. DOI: 10.1038/ismej.2017.16. URL: https://www.nature.com/articles/ismej201716 (visited on 11/21/2025).

[66] Sherwood R. Casjens and Roger W. Hendrix. “Bacteriophage lambda: early pioneer and still relevant”. In: Virology 0 (May 2015), pp. 310–330. ISSN: 0042-6822. DOI: 10.1016/j.virol.2015.02.010. URL: https://pmc.ncbi.nlm.nih.gov/articles/PMC4424060/ (visited on 11/21/2025).

[67] Pavitra Roychoudhury et al. “Fitness benefits of low infectivity in a spatially structured population of bacteriophages”. In: Proceedings of the Royal Society B: Biological Sciences 281.1774 (Jan. 7, 2014). Publisher: Royal Society, p. 20132563. DOI: 10.1098/rspb.2013.2563. URL: https://royalsocietypublishing.org/doi/10.1098/rspb.2013.2563 (visited on 07/09/2025).

[68] Mayra C Obando and Diego O Serra. “Dissecting cell heterogeneities in bacterial biofilms and their implications for antibiotic tolerance”. In: Current Opinion in Microbiology 78 (Apr. 1, 2024), p. 102450. ISSN: 1369–5274. DOI: 10.1016/j.mib.2024.102450. URL: https://www.sciencedirect.com/science/article/pii/S1369527424000262 (visited on 12/08/2025).

[69] Jeanyoung Jo, Alexa Price-Whelan, and Lars E. P. Dietrich. “Gradients and consequences of heterogeneity in biofilms”. In: Nature Reviews Microbiology 20.10 (Oct. 2022). Publisher: Nature Publishing Group, pp. 593–607. ISSN: 1740-1534. DOI: 10.1038/s41579-022-00692-2. URL: https://www.nature.com/articles/s41579-022-00692-2 (visited on 12/08/2025).

[70] Xiaowei Chen et al. “Tide driven microbial dynamics through virus-host interactions in the estuarine ecosystem”. In: Water Research 160 (Sept. 1, 2019), pp. 118–129. ISSN: 0043-1354. DOI: 10.1016/j.watres.2019.05.051. URL: https://www.sciencedirect.com/science/article/pii/S0043135419304397 (visited on 11/20/2025).

[71] Amany M. Al-Anany et al. “Phage Therapy in the Management of Urinary Tract Infections: A Comprehensive Systematic Review”. In: *PHAGE: Therapy*, Applications, and Research 4.3 (Sept. 1, 2023), pp. 112–127. ISSN: 2641-6530. DOI: 10.1089/phage.2023.0024. URL: https://pmc.ncbi.nlm.nih.gov/articles/PMC10523411/ (visited on 11/21/2025).

72. [72] Luisa De Sordi, Marta Lourenço, and Laurent Debarbieux. “The Battle Within: Interactions of Bacteriophages and Bacteria in the Gastrointestinal Tract”. In: Cell Host & Microbe 25.2 (Feb. 13, 2019), pp. 210–218. ISSN: 1931-3128. DOI: 10.1016/j.chom.2019.01.018. URL: https://www.sciencedirect.com/science/article/pii/S1931312819300587 (visited on 11/20/2025).

[73] Marzanna Łusiak-Szelachowska et al. “Bacteriophages in the gastrointestinal tract and their implications”. In: Gut Pathogens 9 (Aug. 10, 2017), p. 44. ISSN: 1757-4749. DOI: 10.1186/s13099-017-0196-7. URL: https://pmc.ncbi.nlm.nih.gov/articles/PMC5553654/ (visited on 11/20/2025).

[74] Désirée A Schmitz et al. “Predicting bacterial interaction outcomes from monoculture growth and supernatant assays”. In: ISME Communications 4.1 (Jan. 1, 2024), ycae045. ISSN: 2730-6151. DOI: 10.1093/ismeco/ycae045. URL: 10.1093/ismeco/ycae045 (visited on 11/21/2025).

[75] Ave T Bisesi et al. “Selection for toxin production in spatially structured environments increases with growth rate”. In: The ISME Journal 19.1 (Jan. 2, 2025), wraf061. ISSN: 1751-7362. DOI: 10.1093/ismejo/wraf061. URL: 10.1093/ismejo/wraf061 (visited on 11/21/2025).

[76] Lin Chao and Bruce R Levin. Structured habitats and the evolution of anticompetitor toxins in bacteria. 1981. DOI: 10.1073/pnas.78.10.6324. URL: https://www.pnas.org/doi/10.1073/pnas.78.10.6324 (visited on 11/25/2025).

[77] Tobias Züst et al. “Using knockout mutants to reveal the growth costs of defensive traits”. In: Proceedings of the Royal Society B: Biological Sciences 278.1718 (Sept. 7, 2011), pp. 2598–2603. ISSN: 0962-8452. DOI: 10.1098/rspb.2010.2475. URL: https://pmc.ncbi.nlm.nih.gov/articles/PMC3136827/ (visited on 11/21/2025).

[78] J. R. Obeso. “The induction of spinescence in European holly leaves by browsing ungulates”. In: Plant Ecology 129.2 (Feb. 1, 1997), pp. 149–156. ISSN: 1573-5052. DOI: 10.1023/A:1009767931817. URL: 10.1023/A:1009767931817 (visited on 11/25/2025).

[79] UniProt. UniProt. URL: https://www.uniprot.org/uniprotkb/Q8LT33/entry (visited on 11/21/2025).

