## Supplementary Information for "Live and let die: lysis time variability and resource limitation shape lytic bacteriophage fitness"

### 1 Single phage killing curve simulation suggests variability in burst size alone has little to no effect on bacteriophage fitness

We ran a preliminary simulation to investigate the effects of variability in burst size and lysis time on a single phage species' ability to clear a bacteria culture. We simulate a protocol which we name the "killing curve": 100 bacteriophages are introduced to a growing bacterial culture, volume $10^{-5}$ml, carrying initially 10,000 cells (carrying capacity 100,000). The infection is allowed to proceed until the number of bacteria drops below 100 (1% of its initial value). We record the time taken for this to occur ($T_{100}$), the maximum count of living bacteria observed ($B_{\text{max}}$), and the number of phages remaining at the end ($V_{\text{end}}$). We then calculate an effective production rate, $(r)$ for the bacteriophage through $r = V_{\text{end}}/T_{100}$.

This simulation is similar to the serial passage simulation discussed in section 3.2 except that only one phage strain was used and there was no transfer to a new culture. The simulation was used to investigate six possible models:

(i) 'All constant', in which $\tau$ and $\beta$ are both completely fixed and deterministic.

(ii) 'Burst size varied', in which $\tau$ is fixed and $\beta$ is drawn from a normal distribution.

(iii) 'Lysis time varied', in which $\tau$ is drawn from a normal distribution and $\beta$ is fixed.

(iv) 'Normal model', in which $\tau$ and $\beta$ are both drawn from uncorrelated normal distributions.

(v) 'Linear model', in which $\tau$ is drawn from a normal distribution and $\beta$ is calculated via $\beta = m'\tau$.

(vi) 'Linear with eclipse model', in which $\tau$ is drawn from a normal distribution and $\beta$ is calculated via $\beta = m(\tau - \varepsilon)$.

Models (iv) through (vi) being those used in the main paper. All parameters used were the same as in serial passage simulations, and can be found in tables 2 to 5. In each case, the simulation was run 20 times and the results were averaged. The simulation was then run an additional 20 times for each model under "conservative" conditions, in which phages did not decay, and only a single phage could bind to each cell.

Figure S1a shows example population dynamics for each burst size model. In all cases the bacterial culture is eradicated in 3 rounds of lysis lasting approximately 17 minutes each. The models which include variable lysis time produce smoother population curves, while the models with deterministic lysis time produce sudden jumps.

The averaged results over 20 simulations (Fig S1b) suggest the following. Variability in lysis time leads to faster bacterial eradication (Fig S1bi). This allows the bacteria less time to divide, reducing the number of available cells (Fig S1bii), and therefore the number of phage produced (Fig S1biii). Hence, variability in lysis time leads to a greater production rate of phages (Fig S1biv) but fewer phages in total.

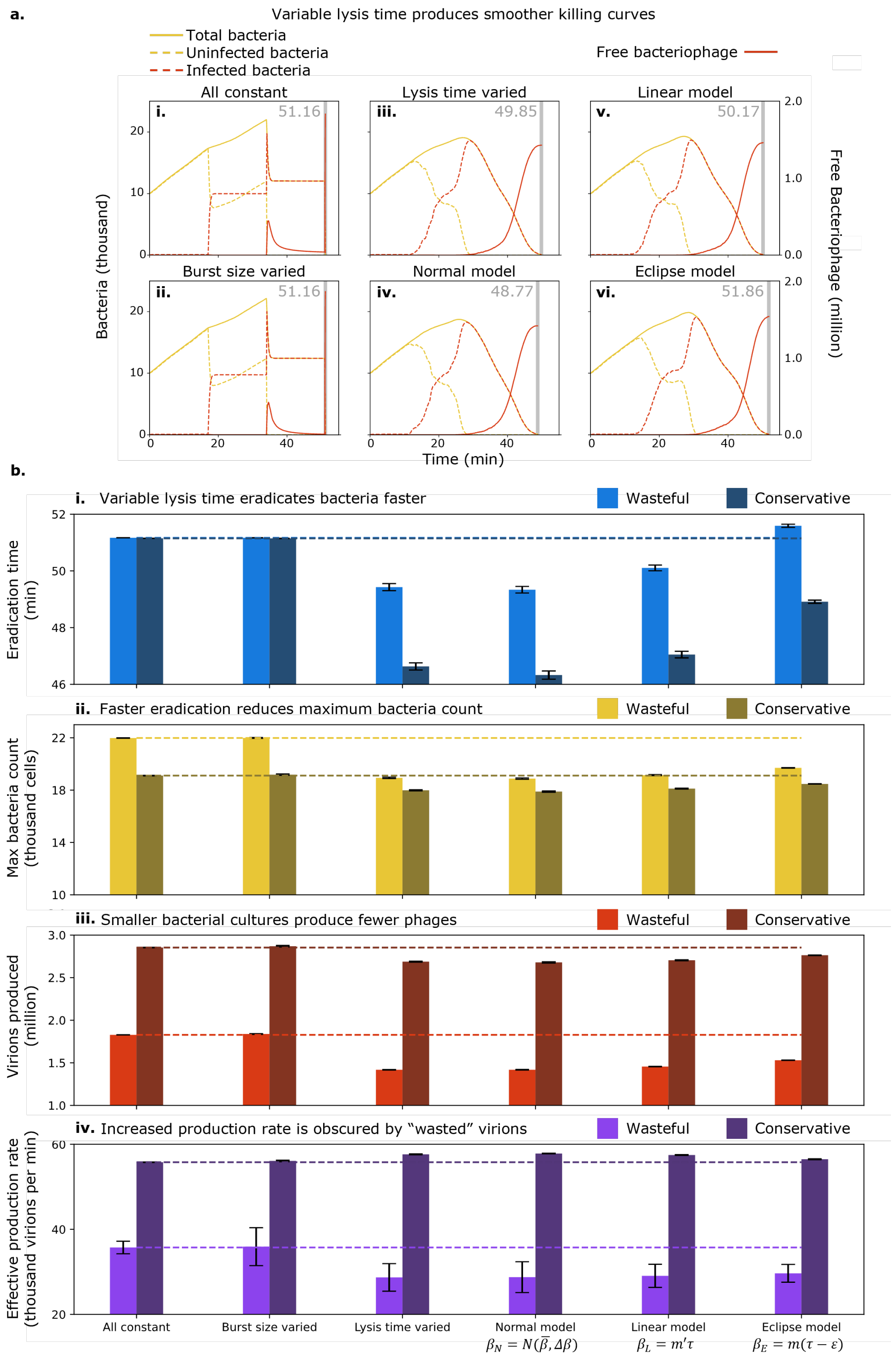


Fig. S1: **a.** Example population curves for the six models tested. The grey vertical line in each panel shows the time at which 99% eradication was achieved and the simulation was terminated. **b.** Results averaged over 20 killing curve simulations showing **i.** Time to 99% eradication,**ii.** Maximum cell count, **iii.** Total virions at time of eradication, **iv.** Virions produced per minute. Error bars show the standard error in the mean.

We also note the significant difference between the "wasteful" and "conservative" cases, and the lack of meaningful difference between the "all constant" and "burst size varied" models, or separately between the "lysis time varied" and "normal" models. While, analytically, we may expect variation in burst size to slightly reduce growth rate (the growth rate $k = \ln(\beta)/\tau$ is concave in $\beta$), under these conditions the MOI quickly becomes much greater than 1 and the laws of exponential growth cease to apply.

In summary, variability in burst size seems to have little to no effect on a phage's ability to eradicate cells or proliferate itself. Lysis time variability however seems to hasten cell culture eradication, at the inevitable cost of fewer virions being produced, as there are fewer cells to consume.

#### 2 Derivation of fitness difference $\Delta F$

Consider two populations, $P_1$ and $P_2$, growing exponentially at rates $k_1$ and $k_2$, such that

$$P_1(t) = P_1(0)e^{k_1 t}$$

and

$$P_2(t) = P_2(0)e^{k_2 t}$$

taking the ratio of these populations at time $t$,

$$\frac{P_2(t)}{P_1(t)} = \frac{P_2(0)}{P_1(0)} e^{(k_2-k_1)t}$$

then taking the natural logarithm of that ratio,

$$\ln\left(\frac{P_2(t)}{P_1(t)}\right) = \ln\left(\frac{P_2(0)}{P_1(0)} e^{(k_2-k_1)t}\right) = \underbrace{\ln\left(\frac{P_2(0)}{P_1(0)}\right)}_{\text{constant}} + (k_2 - k_1)t$$

finally, differentiating with respect to time,

$$\frac{\mathrm{d}}{\mathrm{dt}}\left(\ln\left(\frac{P_2(t)}{P_1(t)}\right)\right) = k_2 - k_1 := \Delta F$$

To avoid ambiguity, we use $k$ to refer to growth rates predicted by numerical integration, and $\Delta F$ to refer to fitness differences measured in the agent based simulation.

If we now consider bacteriophage populations specifically, note that here, $P_i$ must refer to the total population of all virions of type $i$, including those adsorbed to cells. Under the assumptions made, specifically instantaneous adsorption, no super-adsorption, and infinitely abundant bacteria, all phages will be uniquely adsorbed to a cell, hence an equivalent expression for $\Delta F$ is

$$\Delta F_I := \frac{\mathrm{d}}{\mathrm{dt}}\left(\ln\left(\frac{I_2(t)}{I_1(t)}\right)\right).$$

This quantity also works as a measure of fitness difference in resource-limited conditions, because, as one phage grows in population relative to the other and approaches fixation, the ratio of infected cells will converge to 0 or 1.

An alternate expression for $\Delta F$ is

$$\Delta F_V := \frac{\mathrm{d}}{\mathrm{dt}}\left(\ln\left(\frac{V_2(t)}{V_1(t)}\right)\right),$$

which uses the ratio of free virions $V_2/V_1$, as opposed to all virions $P_2/P_1$. This quantity cannot be continuously evaluated as free phage populations frequently fall to zero, so must be evaluated only when it is meaningful, such as just after the transfer step of a serial passage. The $\Delta F_V$ definition does, however, have the advantage that adsorbed phages, which are functionally dead for reproductive purposes, no longer contribute to the fitness. For an analysis of these two possible fitness definitions, see Fig S7. In the main paper we employ $\Delta F_V$ for serial passage simulations, and $\Delta F_I$ for plaque expansions.

### 3 Asymmetric and symmetric lysis time distributions exhibit the same qualitative behaviour

#### 3.1 Erlang distribution

The Erlang distribution is a natural choice for consideration due to its biological relevance: it describes the time taken to complete a series of $n$ exponential processes, each occurring at rate $\lambda$.

$$p(\tau) = \frac{\lambda^k \tau^{k-1} e^{-\lambda\tau}}{(k-1)!} \tag{1}$$

We parametrise our Erlang distribution using the shape $n$ and mean $\overline{\tau} = n/\lambda$. In agreement with Fig 3, under the 'normal' and 'linear' models, for a given mean lysis time, a greater variability (which in the Erlang distribution corresponds to smaller 'shape' $n$) is preferred. Under the 'eclipse' model, a tighter distribution (larger shape) is preferred.

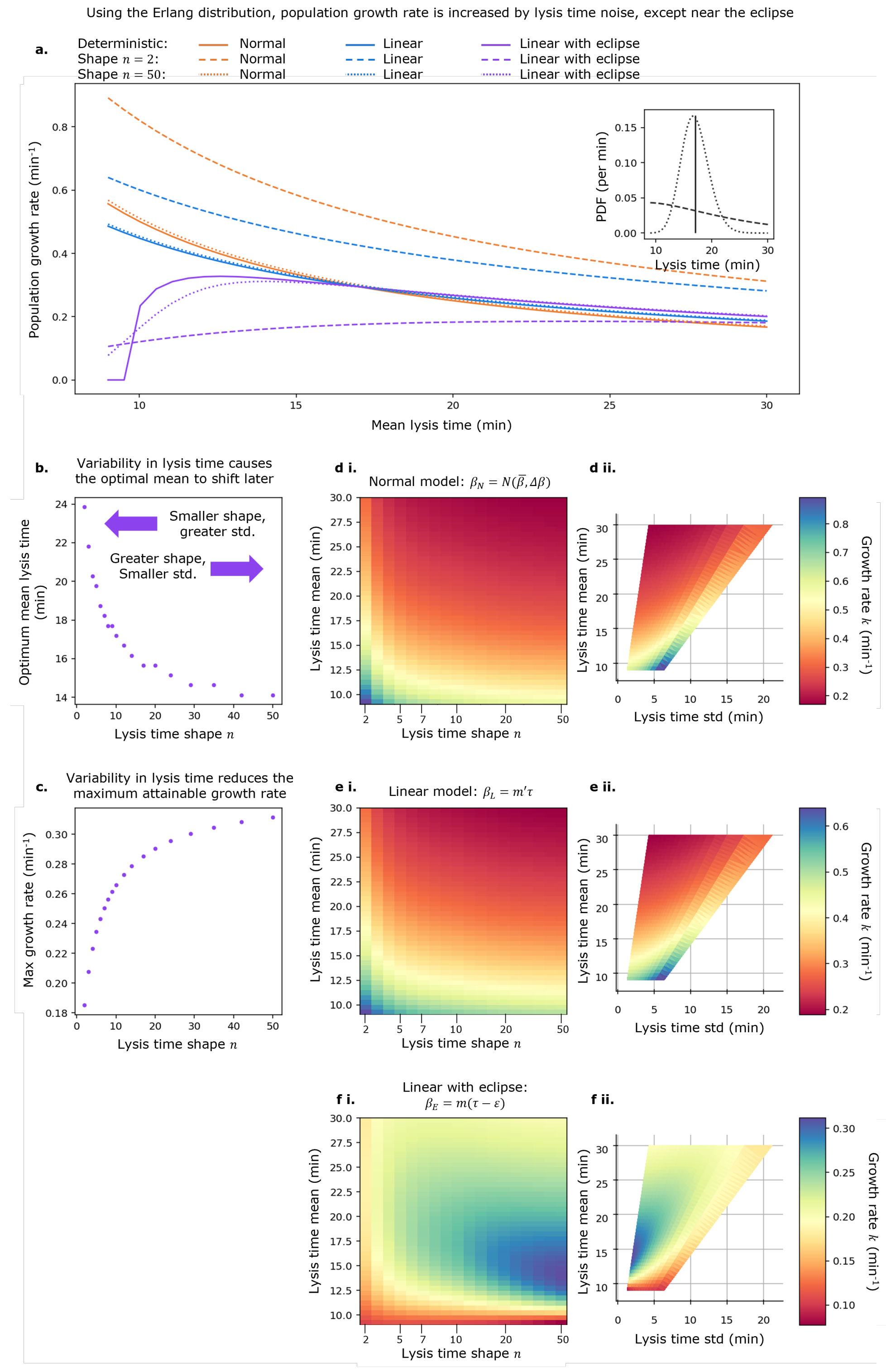


Fig. S2: This figure is a direct comparison to Fig 3.**a.** Bacteriophage effective growth rate $k$ as a function of mean lysis time, for either totally deterministic lysis time (solid lines), Erlang distributed lysis time with shape 2 (dashed lines), or shape 50 (dotted lines). Inset: example lysis time distributions with mean 17 minutes: deterministic (solid line), Erlang with shape 2 (dashed line), Erlang with shape 50 (dotted line). **b.** Optimal lysis time under 'linear with eclipse' model as a function of lysis time shape $n$. **c.** Effective growth rate achieved at optimal lysis time, under 'linear with eclipse' model, as a function of lysis time shape. **d.** to **f.** Effective growth rate as a function of both mean lysis time and lysis time shape $n$ for 'normal', ''linear', and 'linear with eclipse' models respectively (i), and the same data mapped to the lysis time mean and lysis time std axes (ii) to allow easier comparison with Fig 3.

##### 3.2 Weibull distribution

The Weibull distribution usually represents 'time-to-failure', and is described by shape $n$ ($1/n$ is approximately equal to the coefficient of variation) and scale $\lambda$.

$$p(\tau) = \frac{n}{\lambda} \frac{\tau^{n-1}}{\lambda} e^{-(\tau/\lambda)^n} \tag{2}$$

In agreement with Fig. 3, under the 'normal' and 'linear' models, for a given mean lysis time, a greater variability (which in the Weibull distribution corresponds to smaller shape $n$) is preferred. Under the 'eclipse' model, a tighter distribution (larger shape) is preferred.

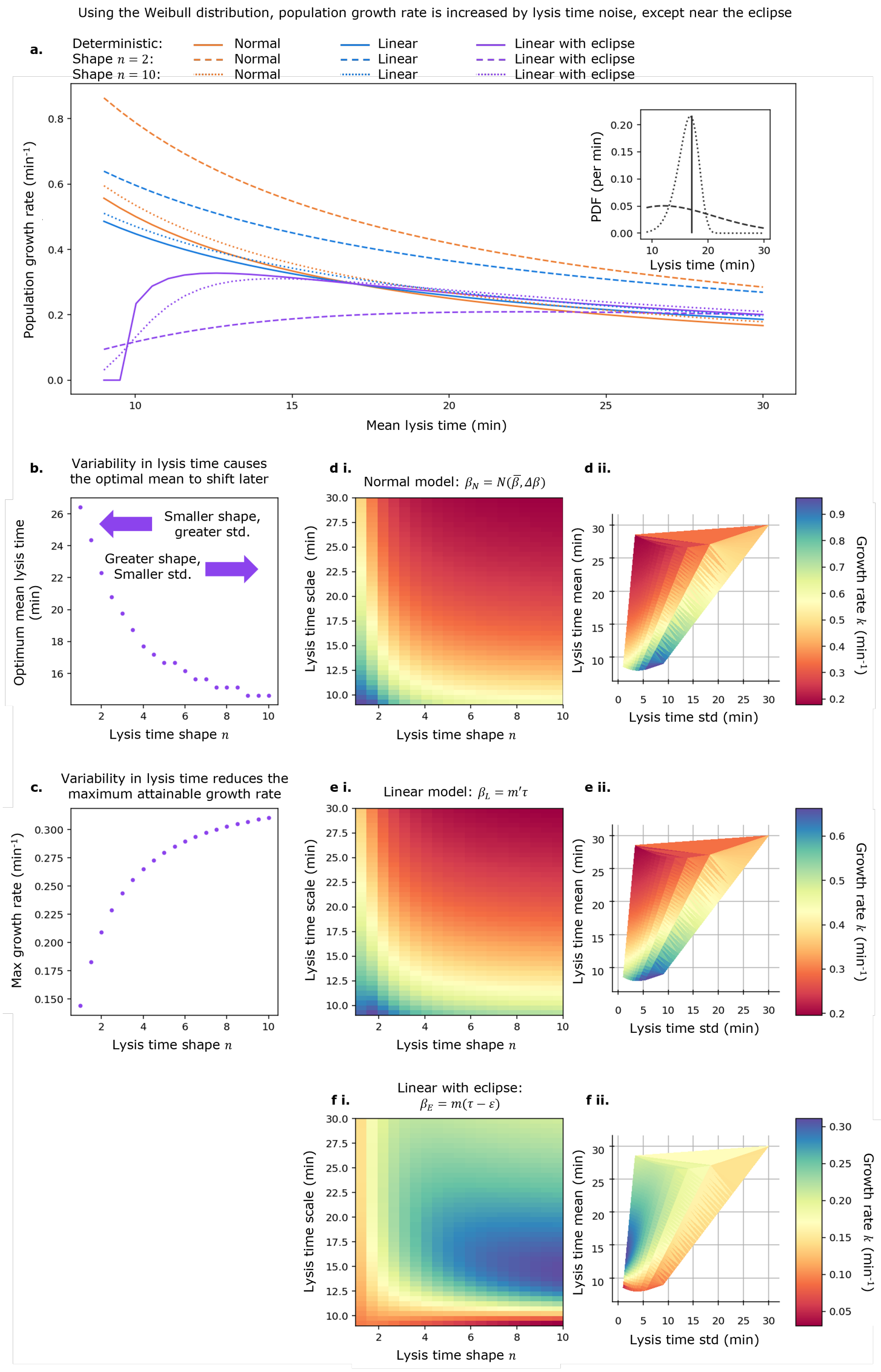


Fig. S3: This figure is a direct comparison to Fig 3. **a.** Bacteriophage effective growth rate $k$ as a function of mean lysis time, for either totally deterministic lysis time (solid lines), Weibull distributed lysis time with shape 2 (dashed lines), or shape 10 (dotted lines). Inset: example lysis time distributions with mean 17 minutes: deterministic (solid line), Weibull with shape 2 (dashed line), Weibull with shape 10 (dotted line). **b.** Optimal lysis time under 'linear with eclipse' model as a function of lysis time shape $n$. **c.** Effective growth rate achieved at optimal lysis time, under 'linear with eclipse' model, as a function of lysis time shape. **d.** to **f.** Effective growth rate as a function of both mean lysis time and lysis time shape $n$ for 'normal', ''linear', and 'linear with eclipse' models respectively (i), and the same data mapped to the lysis time mean and lysis time std axes (ii) to allow easier comparison with Fig 3.

##### 3.3 Skew normal distribution

The skew normal distribution allows us to investigate the effects of skew explicitly. It is described by location $\mu$, size $\sigma$ and skew parameter $\alpha$, and given by:

$$P(\tau) = \frac{2}{\sigma} p_N\big(\frac{x-\mu}{\sigma}\big) \times c_N\big(\alpha \frac{x-\mu}{\sigma}\big), \tag{3}$$

in which $P_N$ and $C_N$ correspond to the standard normal distribution's PDF and CDF respectively.

Defining the quantity $\delta$ as

$$\delta = \frac{\alpha}{\sqrt{1+\alpha^2}},$$

the skew-normal has true mean

$$\overline{\tau} = \mu + \sigma\delta\sqrt{\frac{2}{\pi}},$$

and true variance

$$(\Delta\tau)^2 = \sigma^2\big(1 - \frac{2\delta^2}{\pi}\big).$$

Which can be inverted to define the skew-normal instead through it's true mean, true std and parameter $\alpha$ via

$$\sigma^2 = (\Delta\tau)^2\big(1 - \frac{2\delta^2}{\pi}\big)^{-1},$$

$$\mu = \overline{\tau} - \sigma\delta\sqrt{\frac{2}{\pi}},$$

which is the parametrisation used in this investigation. True skew increases monotonically, though not linearly, with increasing $\alpha$, and is zero at $\alpha = 0$.

We begin by fixing the lysis time std at 2.5 minutes and varying the mean and $\alpha$.

For given mean lysis time, we find that growth rate $k$ depends only weakly on skew. the 'normal' and 'linear' models show increased $k$ for negative skew since this provides a greater likelihood of lysis events happening in near-zero time, which increases effective growth rate. The 'linear with eclipse' model shows a distinct and interesting result: at the lysis time which would be optimal in the totally deterministic case (in which the "distribution" reduces to a delta function) the growth rate obtained is equal for any value of $\alpha$. At lysis times just longer than this optimum positive skew increases growth rate, while just shorter than the optimum negative skew increases growth rate.

Noting (i) that a positively skewed distribution has a steep left face, (ii) that a negatively skewed distribution has a gradually sloping left face and (iii) there is a "cliff edge" in the fitness landscape, in that lysis times below or close to the eclipse period (9.5 minutes) are extremely unproductive, we propose the following: starting with a longer value of lysis time and moving to shorter values, initially the cliff edge can be avoided by utilising a positively skewed distribution with a steep left face, ensuring almost no probability mass is to the left of the cliff edge. As the mean gets closer to the cliff edge, this is no longer possible, so the optimal strategy instead is to switch to a strongly negatively skewed distribution, keeping the peak (mode) to the right of the cliff edge.

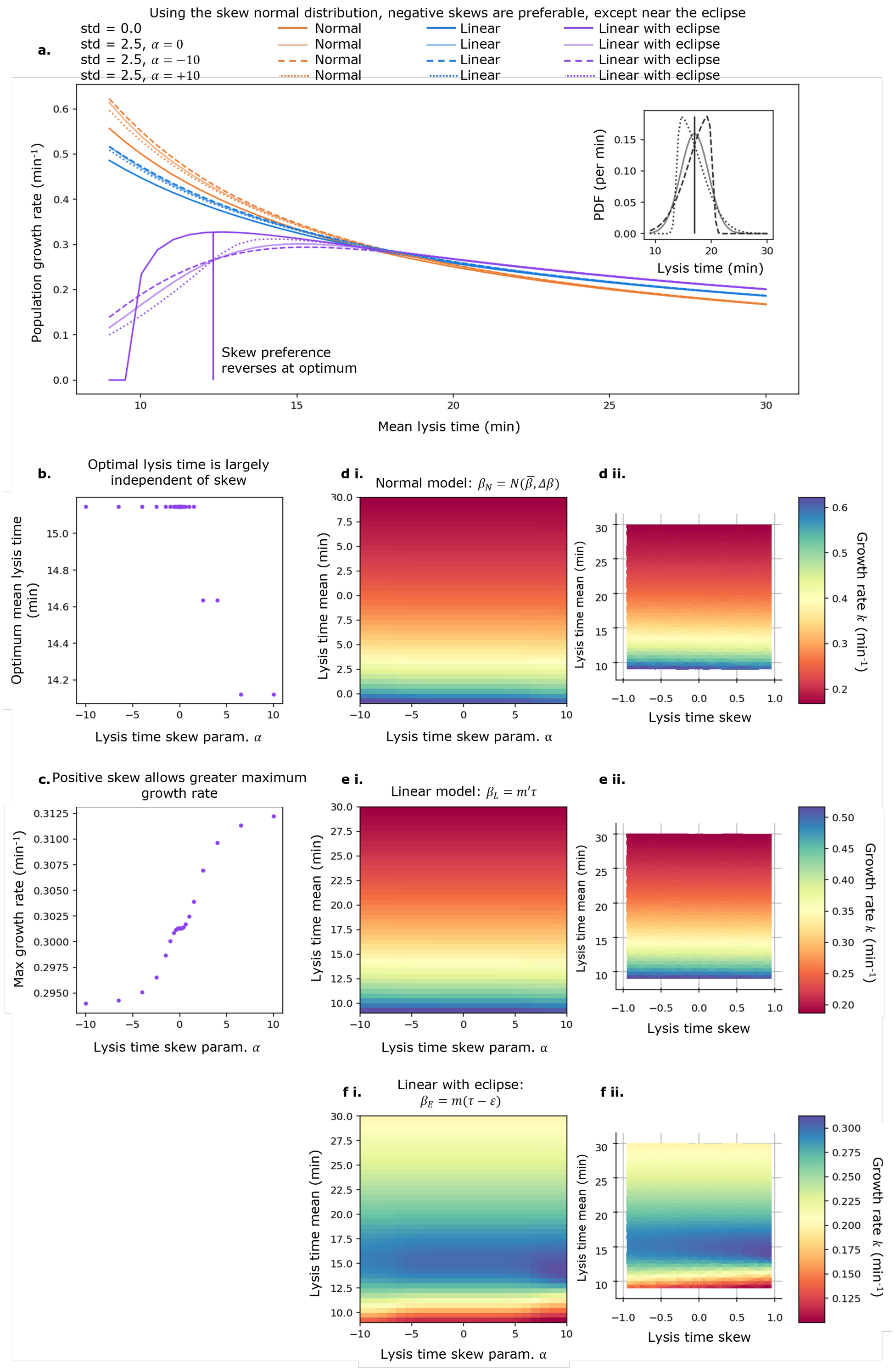


Fig. S4: This figure is a direct comparison to Fig 3. **a.** Bacteriophage effective growth rate $k$ as a function of mean lysis time, for either totally deterministic lysis time (solid lines), skew-normal distributed lysis time with lysis time std = 2.5 min and skew parameter $\alpha = 0$ (pale solid lines), $\alpha = -10$ (strong negative skew, dashed lines), or $\alpha = +10$ (strong positive skew, dotted lines). Inset: example lysis time distributions with mean 17 minutes: deterministic (solid line), skew-normal std = 2.5 min and skew parameter $\alpha = 0$ (pale solid lines), $\alpha = -10$ (dashed lines), or $\alpha = +10$ (dotted lines). **b.** Optimal lysis time under 'linear with eclipse' model as a function of skew parameter $\alpha$. **c.** Effective growth rate achieved at optimal lysis time, under 'linear with eclipse' model, as a function of skew parameter $\alpha$. **d.** to **f.** Effective growth rate as a function of both mean lysis time and skew parameter $\alpha$ for 'normal', ''linear', and 'linear with eclipse' models respectively (i), and the same data mapped to the lysis time mean and true statistical skew axes (ii) to compensate for non-linear relationship between $\alpha$ and true statistical skew.

Next, we fix the mean lysis time at 17 minutes, and vary the std and $\alpha$.

As expected, the 'normal' and 'linear' models show the greatest increase in growth rate for large stds and negative skews. The 'linear with eclipse' model (at the 17 minute mean lysis time chosen) shows the greatest increase for intermediate std and positive skew. This is because 17 minutes is within the convex region of the growth rate function. The curvature,

$$\frac{d^2k}{d\tau^2} = \frac{d^2}{d\tau^2}\left(\frac{\ln(m(\tau-\varepsilon))}{\tau}\right) = \frac{2\ln(m(\tau-\varepsilon)) + \frac{\tau(2\varepsilon - e\tau)}{(\tau-\varepsilon)^2}}{\tau^3},$$

is positive for $\tau \geq 16.75$ minutes.

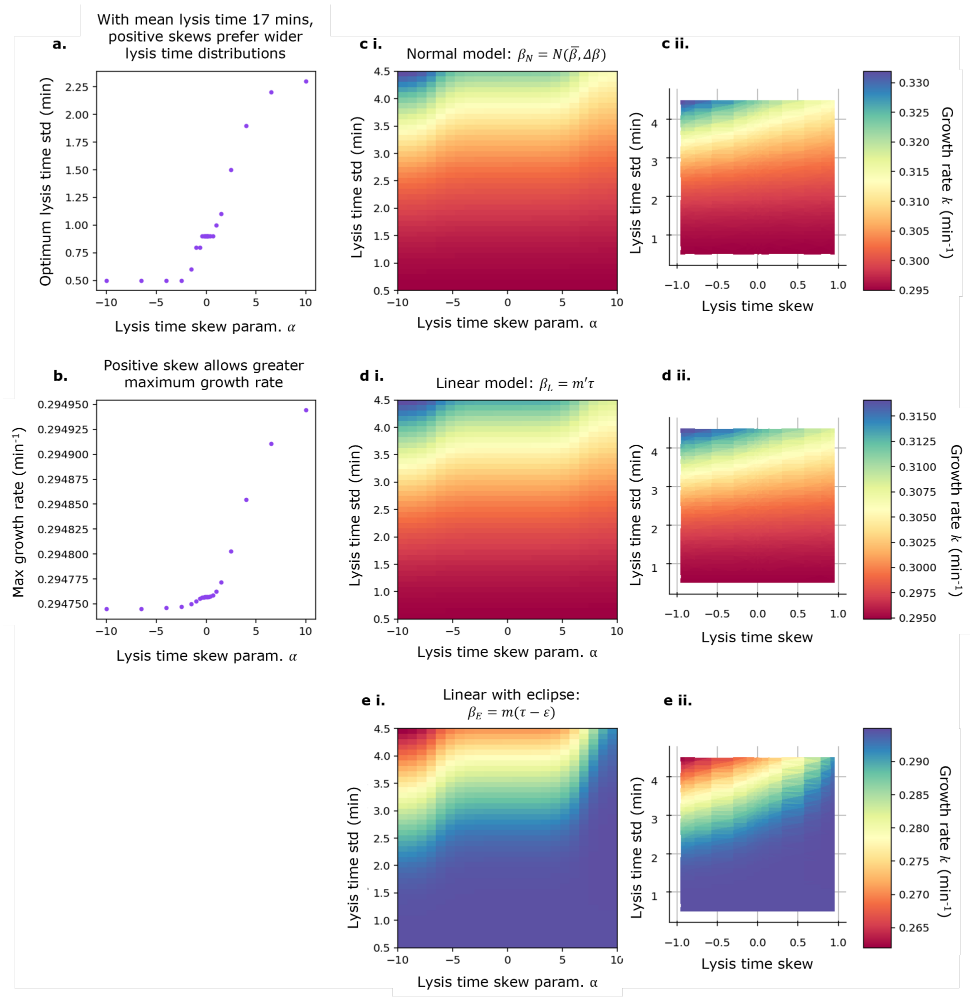


Fig. S5: **a.** Optimal lysis time std under 'linear with eclipse' model as a function of skew parameter $\alpha$. **b.** Effective growth rate achieved at optimal std, under 'linear with eclipse' model, as a function of skew parameter $\alpha$. **c.** to **e.** Effective growth rate as a function of both lysis time std and skew parameter $\alpha$ for 'normal', ''linear', and 'linear with eclipse' models respectively (i), and the same data mapped to the lysis time std and true statistical skew axes (ii) to compensate for non-linear relationship between $\alpha$ and true statistical skew.

### 4 Mean-mean serial passage controls

#### 4.1 Simulation results are independent of absolute scale

All work presented is independent of absolute scale (the absolute number of phages and bacteria considered). Numerically predicted growth rate by default considers only a phage's lysis time $\tau$, burst size $\beta$, and the relationship between them. When corrected to account for adsorption time $\overline{\tau}_{\text{ads}} = 1/\alpha B_0$, dependence is introduced on the adsorption rate $\alpha$ and bacterial density $B_0$. Therefore, the total number of bacteria can be increased by an arbitrary factor and, as long as their concentration is maintained by increasing the volume accordingly, the predicted growth rate is unaffected (Fig S6i).

The serial passage and plaque expansion simulations are also independent of absolute scale, except in the limit of very few virions and cells. If the initial bacteria count, initial phage count, bacterial carrying capacity and simulation volume are all rescaled by the same factor, there is no qualitative or quantitative change in the competition experiment's winner (Fig S6ii), or the winner's fixation rate (Fig S6iii).

Note that, as the absolute scale is reduced (Fig S6 left column), the fixation probability plot (row ii) does appear to "blur" slightly. This is because, at very small initial phage count ($V_0 = 10$), individual rare events can swing the outcome of the simulation significantly, and therefore more variation is seen between the 20 with each set of parameters.

Given this scale-independence, our serial dilution results, computed with 100 phages and 10,000 cells in a volume of $10^{-5}$ ml can be readily applied to a real serial dilution using $10^7$ phages and $10^9$ cells in a volume of 1 ml.

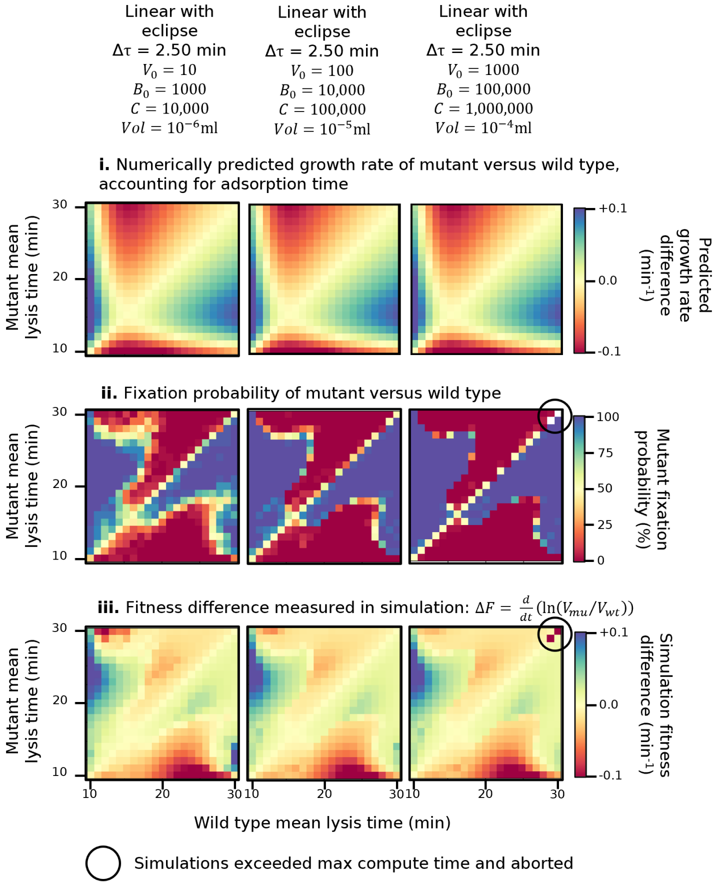


Fig. S6: Results of serial passage competition experiment, competing a wild type phage with one mean lysis time against a mutant with another, at three different absolute scales. **i.** Difference in numerically predicted growth rate, using equation 3. **ii.** Mutant fixation probability in stochastic simulation. **iii.** fitness difference defined as $\Delta F = \frac{\mathrm{d}}{\mathrm{dt}}\left(\ln\left(\frac{V_{\mathrm{mu}}}{V_{\mathrm{wt}}}\right)\right)$. The middle column here is identical to the rightmost column of Fig 4.

##### 4.2 Fitness difference can be calculated through virion abundance or number of cells infected

In section 2, we describe how the fitness difference in simulation $\varDelta F$ can be calculated using either the total free phage

$$\varDelta F_V = \frac{\mathrm{d}}{\mathrm{dt}}\left(\ln\left(\frac{V_{\mathrm{mu}}}{V_{\mathrm{wt}}}\right)\right),$$

derived in 2, or using the number of infected cells

$$\varDelta F_I = \frac{\mathrm{d}}{\mathrm{dt}}\left(\ln\left(\frac{I_{\mathrm{mu}}}{I_{\mathrm{wt}}}\right)\right).$$

In the main paper, we use the $\varDelta F_V$ definition for serial passage simulations. In plaque expansion simulations, however, the 'treadmill' operation removes virions from the back of the travelling infection wave, so the quantities $V_{\mathrm{wt}}$ and $V_{\mathrm{mu}}$ are not meaningful away from the wavefront. We therefore use the $\varDelta F_I$ definition.

To compare these definitions of $\varDelta F$, we present both here for the serial passage experiment described in Fig 4. In serial passage, both quantities are well defined, though not necessarily equivalent, if measured just before the transfer step. $V_{\mathrm{mu}}$ is the number of mutant virions remaining at time of transfer, and $I_{\mathrm{mu}}$ is the total number of cells infected by the mutant in the current passage cycle.

We see that the two definitions broadly agree, except that some features on the $\varDelta F_V$ plots are not captured by the the $\varDelta F_I$ plots (Fig S7 iii and iv). $\varDelta F_V$ seems to be the better metric, in that it everywhere agrees with the mutant fixation probability. Disagreements seem mostly localised to the fitness "stripes", and are most pronounced under the 'linear' and 'linear with eclipse' models. The rationale for this is as follows. First, stripes represent regions where adsorption and decay shape the fitness landscape more than growth rate, hence $\varDelta F_I$ may predict a phage should win as it infected more cells, but $\varDelta F_V$ may capture that many of those virions then decayed. Second, these stripes are some distance from the line $y = x$, so, under the 'linear' and 'linear with eclipse' models, in which the stripes are more prominent, the disparity in lysis times and therefore burst sizes means there may be a significant difference in the ratios $V_{\mathrm{mu}}/V_{\mathrm{wt}}$ and $I_{\mathrm{mu}}/I_{\mathrm{wt}}$.

Because $\varDelta F_V$ captures more features of the fitness landscape and better agrees with mutant fixation probability, this is the metric used in serial passage simulations.

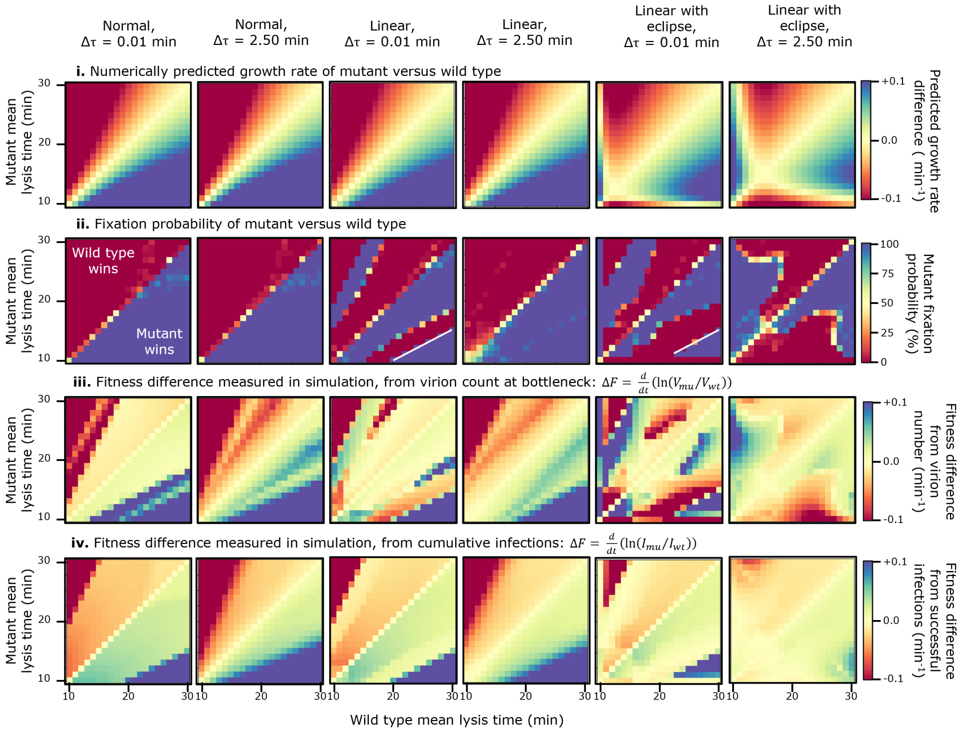


Fig. S7: This figure is a direct comparison to Fig 4. Results of serial passage competition experiment, competing a wild type phage with one mean lysis time against a mutant with another. **i.** Difference in numerically predicted growth rate, using equation 3. **ii.** Mutant fixation probability in stochastic simulation. **iii.** fitness difference defined as $\Delta F_V = \frac{\mathrm{d}}{\mathrm{dt}}\left(\ln\left(\frac{V_{\mathrm{mu}}}{V_{\mathrm{wt}}}\right)\right)$. **iv.** fitness difference defined as $\Delta F_I = \frac{\mathrm{d}}{\mathrm{dt}}\left(\ln\left(\frac{I_{\mathrm{mu}}}{I_{\mathrm{wt}}}\right)\right)$.. Rows i to iii are identical to those in Fig 4.

#### 4.3 Population resonance creates fitness "stripes"

To demonstrate the mechanism that gives rise to the observed "stripes" on the serial passage fitness landscape, we present example population traces from within and just to the right of the stripe for each of the three burst size models Fig S8a. In all cases the mutant phage has a mean lysis time of $\overline{\tau}$ of 12 minutes, the wild type has mean $\overline{\tau}$ 23 (i, iii, v) or 25 minutes (ii, iv, vi), and both have standard deviation $\Delta\tau = 0.01$ minutes.

By inspection of the population traces, we see that, within the stripe, the wild type delays fixation of the mutant (i) or reaches fixation itself (iii, v), by lysing later, infecting a large fraction of the available cells, and forcing the mutant to endure a significant delay between completing its final round of lysis and undergoing a passage, during which time is suffers virion loss due to super-adsorption and decay. If the mutant's mean lysis time is less than half that of the wild type (ii, iv, vi), this strategy is no longer viable, as the mutant's second round of lysis now secures almost all of the available resources.

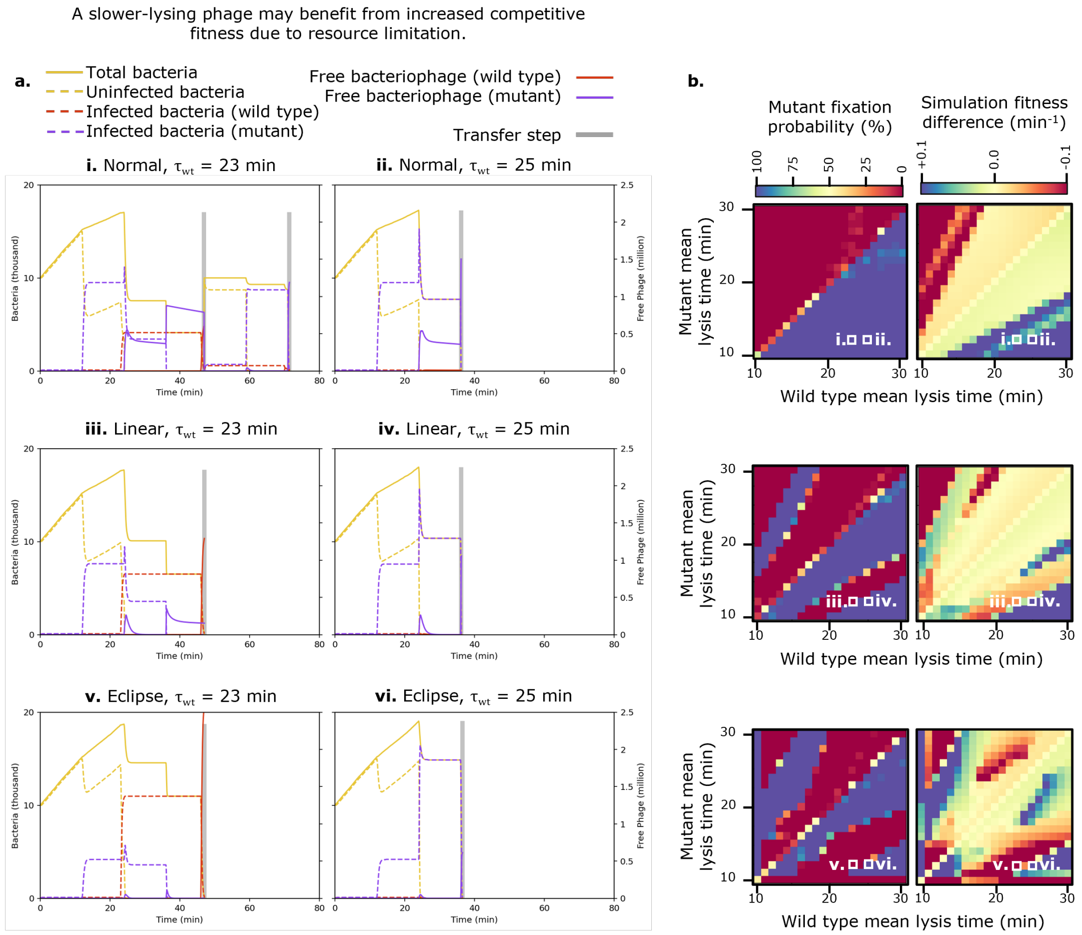


Fig. S8: **a.** Population vs time plots for all three burst size models to illustrate fitness "stripes". In all competitions, the mutant page had a mean lysis time of 12 minutes, and both phages had a lysis time std of 0.01 minutes. **b.** Copy of the data presented in Fig. 4: mutant fixation probability (left column) and fixation rate (right column) as a function of mutant and wild type mean lysis time. Parameter values corresponding to the traces on the left have been highlighted.

##### 4.4 Serial passage transfer interval creates additional resonance effects

When conducting a serial passage experiment, one may choose to perform transfers at fixed time intervals [1, 2] or after the bacterial culture clears [3]. As discussed in section 3.2, if a periodic transfer is used, and the period is long enough that all or almost all bacteria have been lysed, then the only difference compared to transfer-on-clearing is how long free virions are left to decay in solution. If, however, the transfer occurs before all cells have been lysed, additional resonant effects are introduced by the transfer period (Fig S9).

We repeat the simulations described in Fig 4, but now the 1000X dilution occurs every 25 minutes. The simulation runs for 150 minutes (6 transfers) and then terminates. Results from the original transfer-on-clearing simulation are presented in Fig S9i, while the results of the 25 minute periodic transfer are presented in Fig S9ii. As the dilution was now transferring infected cells alongside (and often instead of) free virions, we calculate fitness difference based on the number of successful infections per passage, rather than the number of free virions immediately following a transfer. See figure S7 for a comparison of these two definitions.

This definition of $\Delta F$ tends to somewhat mask the population resonance effect. Even so, a sharp fitness discontinuity at $y = x/2$ can be seen in all columns of Fig S9i and the first two columns of Fig S9ii. This corresponds to the sudden gain in fitness a mutant experiences by setting its mean lysis time less than half that of the wild type, thereby allowing an additional round of lysis.

In the leftmost plot of row ii, corresponding to the 'normal' relation between burst size and lysis time, we see three new fitness discontinuities compared to row i. These discontinuities are located at $y = T$, $y = 2T/3$, and $y = T/2$ for $T = 25$ min. The greatest discontinuity is located at $y = T$, which corresponds to the fact that a mutant with lysis time greater 25 minutes cannot amplify fast enough to overcome the 25-minute periodic dilution, and will quickly be lost. Interestingly, the sign of the other discontinuities suggests that a lysis time longer than $T/2$ but shorter than $2T/3$ is advantageous. The benefit of having a lysis time shorter than $2T/3$ is straightforward: a mutant in this region will achieve three lysis cycles per two passages, relying on the fact that infected cells as well as free phages are transferred. The benefit of having a lysis time longer than $T/2$ is less intuitive, especially since these results were obtained using the 'normal' model, so there is no benefit to longer lysis cycles in terms of virions produced.

This discontinuity arises from the population resonance effect as follows: there are enough susceptible cells available to support two lysis cycles per phage pool. By the end of the second cycle, there are few cells remaining, so a third cycle would be attempted at very high MOI with many virions lost to super-adsorption. Additionally, for a mutant with lysis time just shorter than $T/2$, shortly after this wasteful third cycle begins, a transfer occurs, supplying many new cells which are inaccessible to the phage until its third lysis cycle completes. By contrast, a mutant with lysis time just greater than $T/2$ will, at the time of the transfer, have infected a large number of cells, some of which will survive the dilution and then burst almost immediately, releasing their virions into a fresh pool of susceptible bacteria.

The other two plots in row ii are more complex, as now a phage's fitness depends on its lysis time through three distinct mechanisms: its effect on burst size, resonance with the lysis time of the competing phage, and resonance with the transfer period. That the fitness landscapes here show similar features to those in Fig 4 but are harder to interpret motivates our decision to simulate transfer-on-clearing.

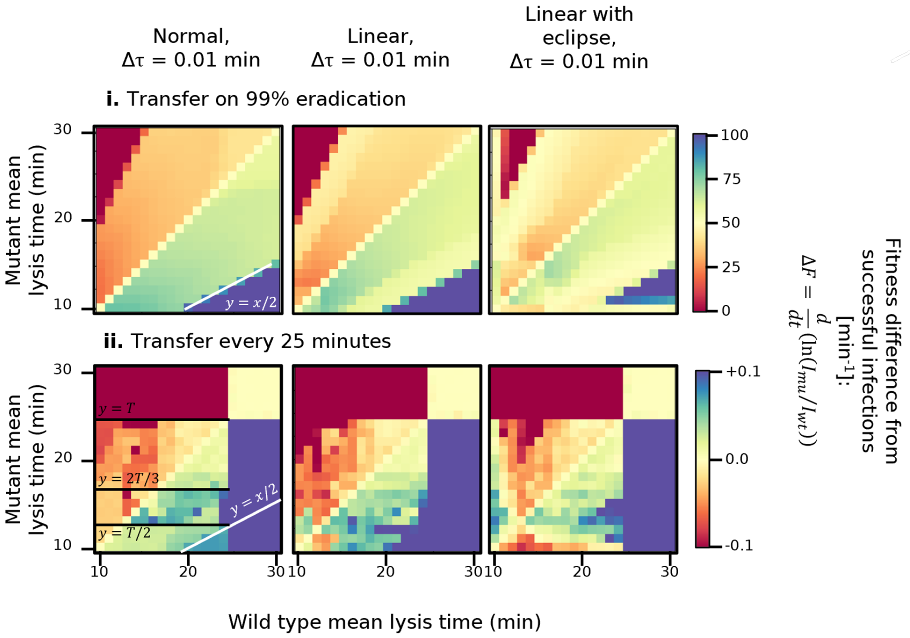


Fig. S9: This figure is a direct comparison to Fig 4. Results of serial passage competition experiment, competing a wild type phage with one mean lysis time against a mutant with another. **a.** Same data is Fig 4, super-adsorption, free virion decay, and cell growth enabled ($A_{\max} = 100, \delta = 0.01, B_0 = 10,000, C = 100,000$). **b.** Super-adsorption and free virion decay disabled, cell growth enabled ($A_{\max} = 1, \delta = 0, B_0 = 10,000, C = 100,000$). **c.** Super-adsorption, free virion decay and cell growth disabled ($A_{\max} = 1, \delta = 0, B_0 = 10,000, C = 10,000$). Top row in a. is growth rate difference predicted by numerical integration, which assumes no super-adsorption or decay and abundant bacteria. Middle row in a. and top rows in b. and c. are mutant fixation probability. bottom rows in a. b. and c. are simulation fitness difference $\Delta F$. In all rows apart from fixation probability the visual scale has been set to saturate at $\pm 0.1$ min$^{-1}$.

#### 4.5 Population resonance requires super-adsorption and decay

To further demonstrate that the ”stripes” on the serial passage fitness landscape arise from virion loss due to super adsorption and decay, we repeat all simulations discussed in section **??** (serial passage competition of two phage strains with different mean lysis times) under different conditions. In Fig S10a we reproduce the results of figure 4 without modification. In Fig S10b, we remove the possibility of virion loss by disabling both super-adsorption and decay ($A_{\mathrm{max}} = 1, \delta = 0$). Aside from the step fitness increase at $y = x/2$, agreement between the measured and predicted fitness differences is improved. When we further disable cell growth (Fig S10c) agreement improves further, though is still not complete.

A persistent distinction between predicted and measured fitness differences is that, in the serial passage case, because resources are finite, a number of step discontinuities in fitness will always be present, especially when lysis time std is small, since completing a lysis cycle even only slightly faster than an opponent is rewarded by a great share of resources.

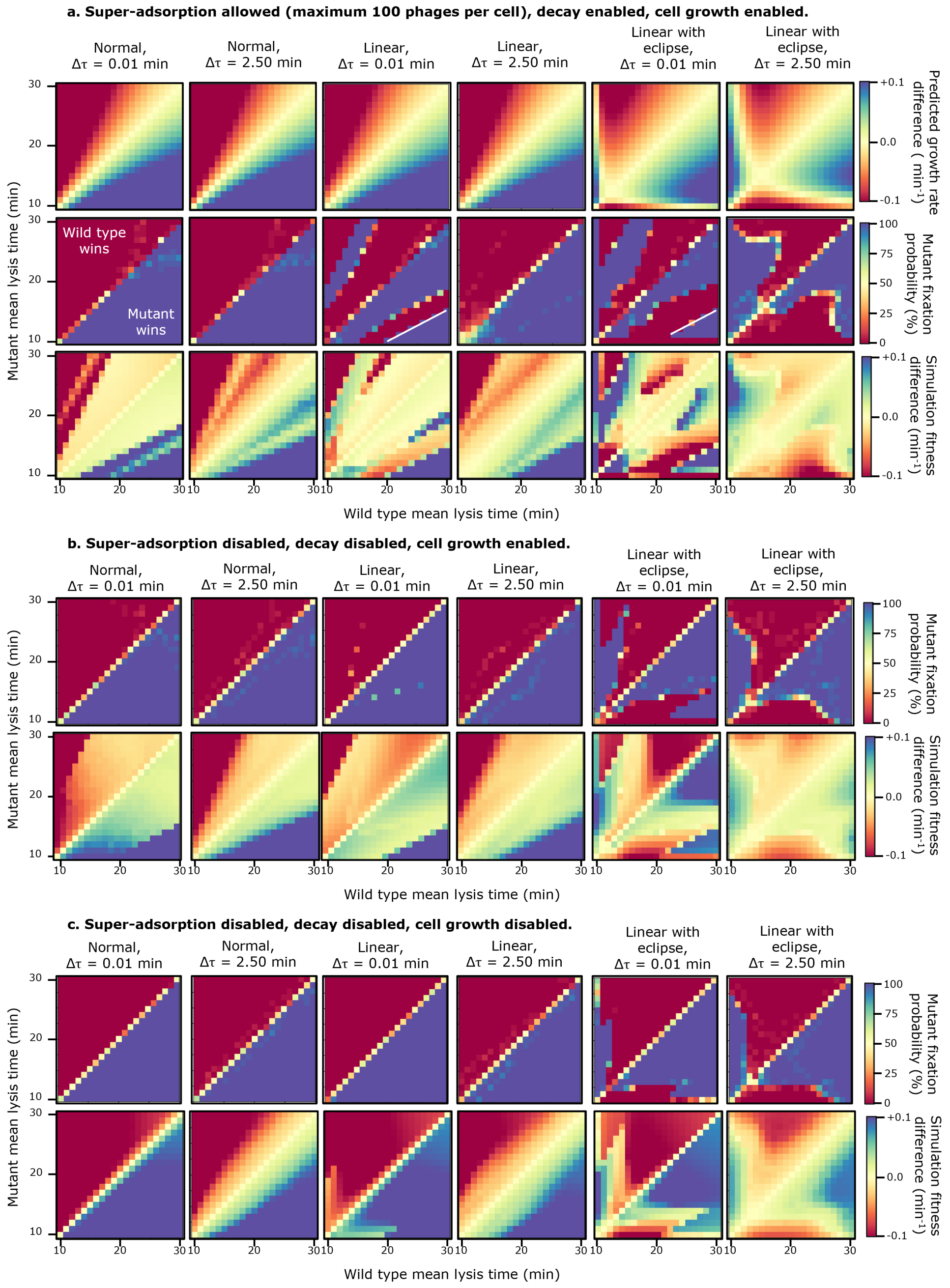


Fig. S10: This figure is a direct comparison to Fig 4. Results of serial passage competition experiment, competing a wild type phage with one mean lysis time against a mutant with another. **a.** Same data is Fig 4, super-adsorption, free virion decay, and cell growth enabled ($A_{\text{max}} = 100, \delta = 0.01, B_0 = 10,000, C = 100,000$). **b.** Super-adsorption and free virion decay disabled, cell growth enabled ($A_{\text{max}} = 1, \delta = 0, B_0 = 10,000, C = 100,000$). **c.** Super-adsorption, free virion decay and cell growth disabled ($A_{\text{max}} = 1, \delta = 0, B_0 = 10,000, C = 10,000$). Top row in a. is growth rate difference predicted by numerical integration, which assumes no super-adsorption or decay and abundant bacteria. Middle row in a. and top rows in b. and c. are mutant fixation probability. bottom rows in a. b. and c. are simulation fitness difference $\Delta F$. In all rows apart from fixation probability the visual scale has been set to saturate at $\pm 0.1$ min$^{-1}$.

##### 4.6 Slower adsorption desynchronises lysis cycles, removing resonance stripes.

If uninfected bacterial hosts are present at low but constant density $B$, and the phage has adsorption rate $\alpha$, the "adsorption time" $\tau_{\text{ads}}$ (time taken for a virion to find and bind to bacterial cell) will be exponentially distributed with mean $\overline{\tau}_{\text{ads}} = 1/\alpha B$.

$$p_{\text{ads}}(\tau_{\text{ads}}) = \alpha B e^{-\alpha B t} \tag{4}$$

Generation times $T$ are now composed of lysis time $\tau_{\text{lys}}$ and adsorption time $\tau_{\text{ads}}$ through $T = \tau_{\text{lys}} + \tau_{\text{ads}}$. The distribution of generation times $p(T)$, is thus given by the convolution

$$p_{\text{tot}}(T) = p_{\text{lys}}(\tau_{\text{lys}}) * p_{\text{ads}}(\tau_{\text{ads}}) = \int_0^T p_{\text{lys}}(t) p_{\text{ads}}(T-t) \mathrm{d}t. \tag{5}$$

The upper limit of integration being $T$ as $p_{\text{ads}}(T-t)$ is undefined for $(T-t) \leq 0$. The growth rate $k$ is then

$$k = \int_{T=0}^{\infty} \int_{t=0}^{T} \frac{p_{\text{lys}}(t) p_{\text{ads}}(T-t) \ln \beta(t)}{T} \mathrm{d}t \mathrm{d}T. \tag{6}$$

The adsorption rate $\alpha$, initial bacterial number $B_0$ and simulated volume *Vol* used in the serial passage experiments discussed in section **??** and Fig 4 describe a system quite close to the high-adsorption limit ($\overline{\tau}_{\text{ads}} = 20$ seconds). To investigate the sensitivity of our results to this limit, we repeated the serial passage assay using the same parameters, except that the simulated volume *Vol* was increased by factor 100, representing a much sparser bacterial culture (Fig S11 without increasing the initial number of bacteria or the carrying capacity. This is mathematically equivalent to a reduction in adsorption rate.

In all three burst size models, numerically predicted growth rate predicts a global reduction in fitness difference $k_{\text{mu}} - k_{\text{wt}}$, across the whole of parameter space, since all phages will reproduce more slowly when the time to locate a host increases. The optimal lysis time predicted by the 'linear with eclipse' model also increases. As resources are harder to find, the explore-exploit trade-off shifts in favour of spending longer infecting each cell.

Simulated fitness difference $\Delta F$ and mutant fixation probability largely agree, with two interesting points to note. Firstly, the desynchronisation of lysis cycles caused by the adsorption delay removes any population resonance stripes. Second, the optimal lysis time for the 'linear with eclipse' model, shifts more than expected. Growth rate maximisation without considering adsorption time places the optimum at 15 minutes. Considering adsorption time places the optimum at about 17 minutes, and the simulation places the optimum at about 20 minutes. We interpret this as arising from resource limitation. Even though adsorption is now slow enough that super-adsorption is less of a concern, the limited nature of the available resources still rewards efficiency (taking longer to lyse but producing more offspring per cell).

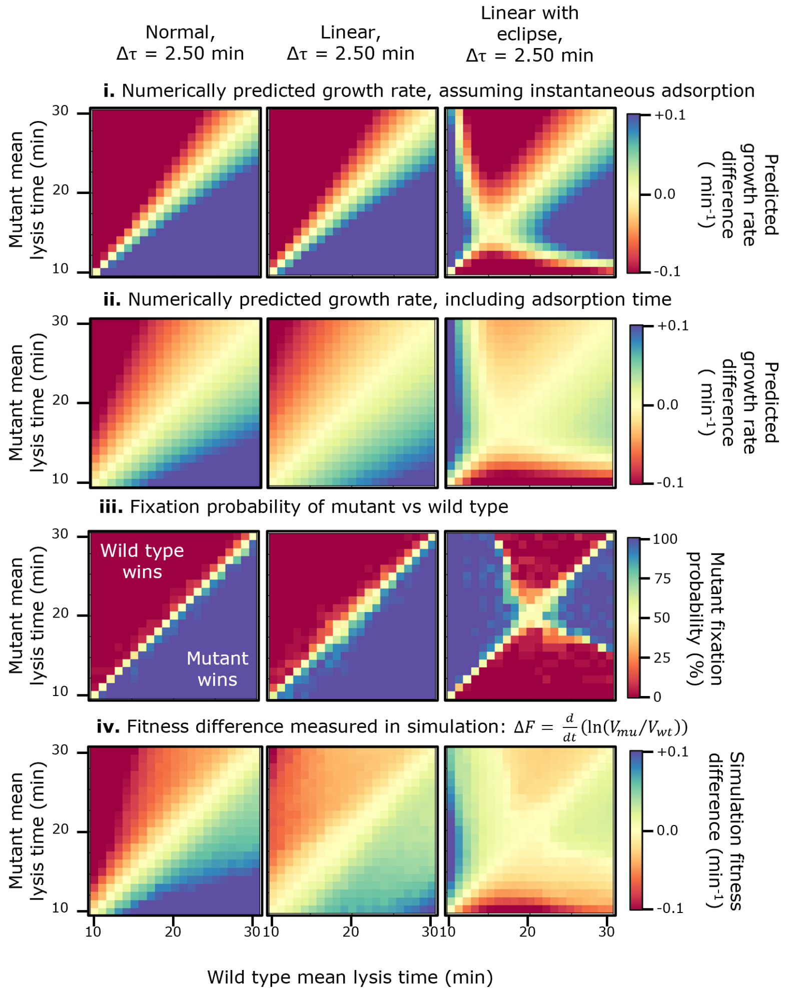


Fig. S11: This figure is a direct comparison to Fig 4. Results of serial passage competition experiment, competing a wild type phage with one mean lysis time against a mutant with another, in a low-adsorption regime. **i.** Difference in numerically predicted growth rate, using equation 3 without correcting for adsorption time. **ii.** Difference in numerically predicted growth rate, using equation 3 convolved with an exponential distribution of mean $1/\alpha B_0$ to correct for adsorption time. **iii.** Mutant fixation probability in stochastic simulation. **iv.** Fitness difference in stochastic simulation.

### 5 Mean-mean minimal simulation controls

#### 5.1 Qualitative observations are robust to the choice of phage parameters

To confirm that our observations are not unique features of our specific parameter choices, we investigate a number of additional parameter choices using both numerical integration and a minimal "fast" simulation 5.3. First, we note that in our T7-*E. coli* inspired system, the range of lysis times available (10 to 30 minutes) is of similar order to the bacterial doubling time (20 minutes).

Changing the lysis time necessarily changes the burst size, so in order to investigate this relationship between lysis time and doubling time without confounding variables, we consider two phages in competition, predating bacteria with doubling times of 4, 9, 20 and 100 minutes (Fig S12).

We note that doubling times much greater than the lysis time do not introduce significantly and qualitatively new behaviour, but that doubling times much faster than lysis create a more complex fitness landscape. This is because, with bacteria doubling every 4 minutes (Fig S12vi), even a very small change in lysis time drastically alters the number of available cells. There is also an increase in the numerical instability of the 'fast' simulation as the bacterial doubling time decreases, visible as a yellow "speckle" across the fitness landscape.

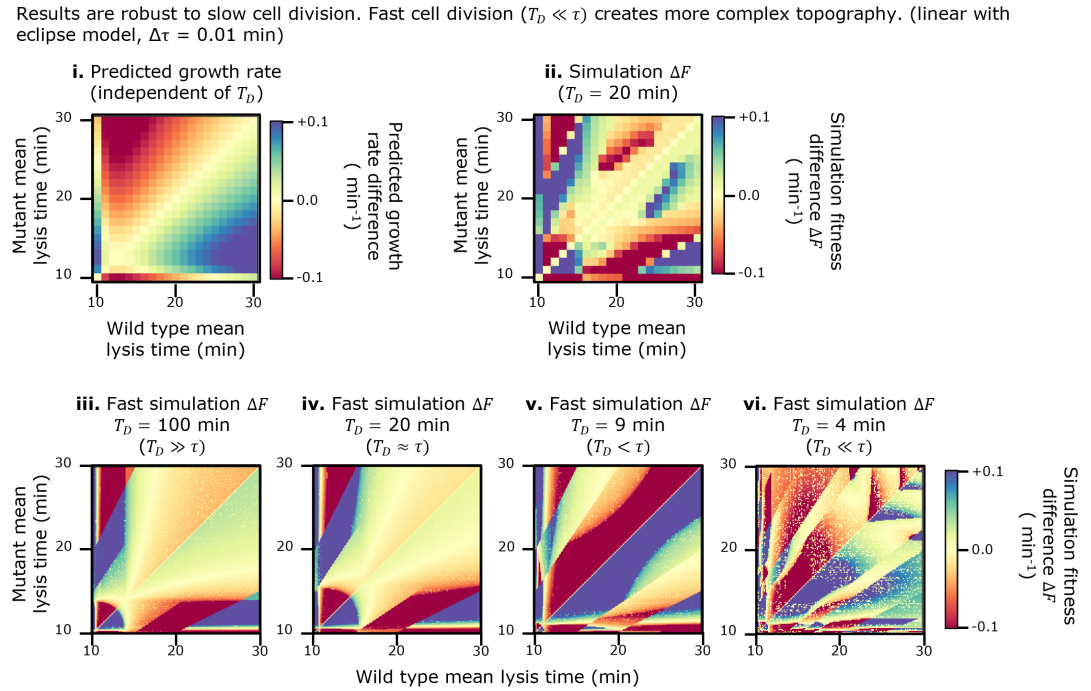


Fig. S12: **i.** Predicted growth rate difference between two phages in competition using the 'linear with eclipse' model, 0.01 min lysis time std. This prediction is made without reference to any bacterial doubling time, as it assumes bacteria are abundant. **ii.** Fitness difference of the same phages in serial passage simulation. Panels i and ii both appear in Fig 4, in rows i and iii respectively. **iii** to **vi**. Fast simulation results showing the effect of reducing bacterial doubling time. Panel iv corresponds to the same parameter values as panel ii, but using the 'fast' simulation 5.3 rather than the full simulation.

Next we note that burst sizes for different phage-bacteria pairs vary over several orders of magnitude [4]. To investigate the sensitivity of our results to our choice of burst sizes (centred on 150), we again consider two phages in competition, using the 'linear with eclipse' model, and vary either the maturation rate or eclipse period (Fig S13).

Numerical growth rate predictions (Fig S13a) suggest that varying $m$ and $\varepsilon$ alters the optimal lysis time, and the steepness of the iso-fitness contours, but does not introduce any new qualitative results. The 'fast' simulation suggests that very low maturation rates create a complex and unstable fitness topography, due to the phage propagating very slowly, and thus the relative growth rate of the bacteria being very large.

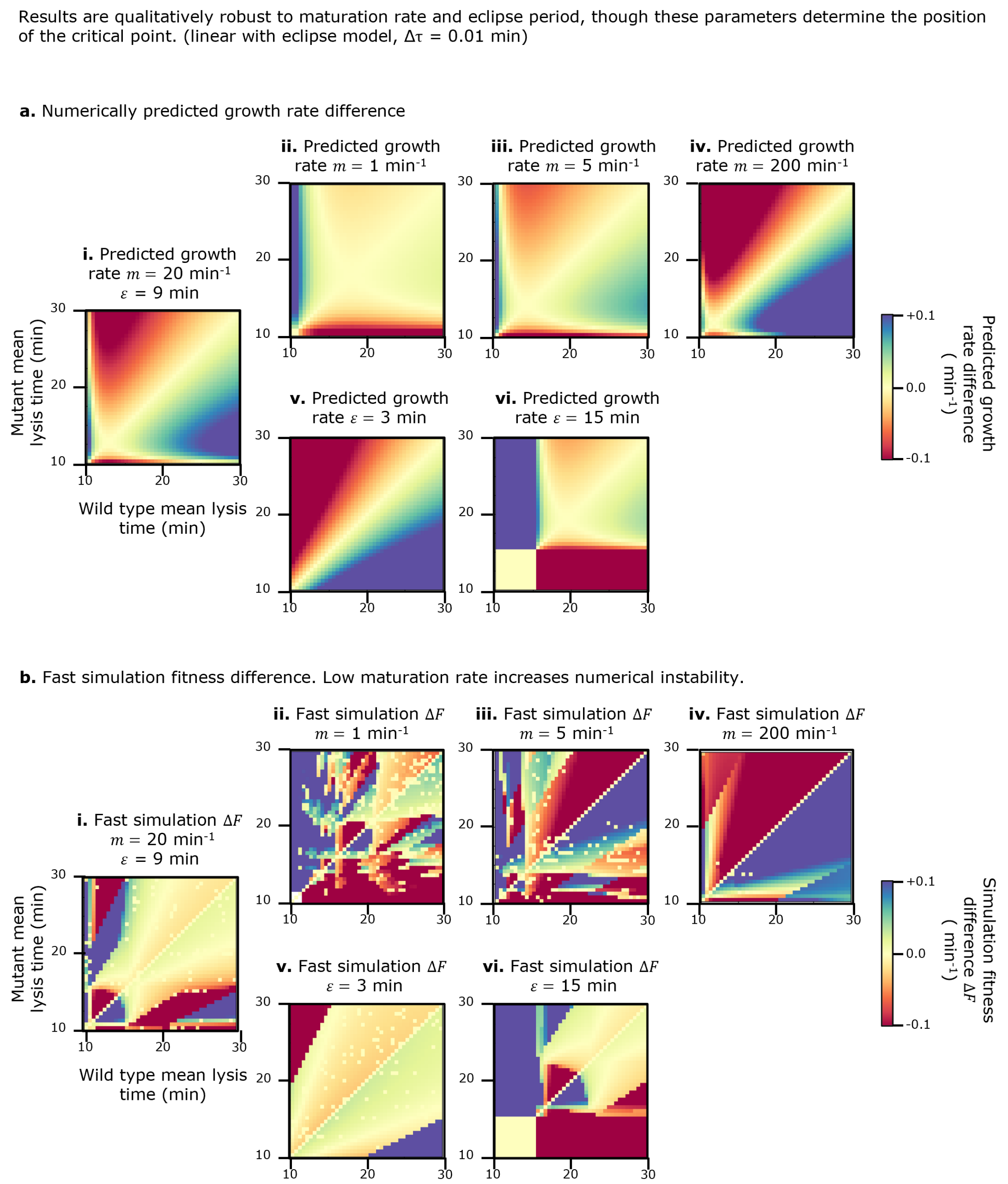


Fig. S13: **a.** Predicted growth rate difference between two phages in competition using the 'linear with eclipse' model, 0.01 min lysis time std, for the parameters used in the main paper (i), for varied maturation rates $m$ (ii to iv) and varied eclipse periods $\varepsilon$ (v to vi). **b.** Same as a. but showing 'fast' simulation fitness difference.

#### 5.2 Population resonance depends on absolute cell count

To investigate the role of initial cell count in the population resonance phenomenon, we employ a fast-running "minimal" simulation. This simulation makes two additional assumptions: (i) that lysis times are totally deterministic, and (ii) that adsorption is instantaneous. This means all cells infected by a phage strain (mutant or wild type) will lyse at once, and the virions released will then immediately adsorb, up to the limit of maximum adsorbed virions per remaining cell. In figure S14, we plot the fitness difference $\Delta F$ between the mutant and wild type phage as their mean lysis times are varied from 1 to 50 minutes, for a range of initial cell numbers $B_0$. The 'normal' model was employed in this analysis, both phages using a common burst size of 150.

We note two distinct changes in the topography of the fitness landscape as the number of susceptible cells is increased, as well as a degree of numerical noise which is most prevalent when cell count is small (Fig S14i,ii). Since these plots are antisymmetric by construction, we will again limit discussion to the bottom right hand side of the parameter space, below the line $y = x$, without loss of generality.

First, we see a number of discontinuous changes in $\Delta F$, each aligned with a particular ratio $y/x$ (labelled explicitly in Fig S14xi). The most prominent of these is $y/x = 1/2$, which is visible to a greater or lesser extent in all panels. As the cell count is increased fitness discontinuities at $y/x = 1/3$, $y/x = 2/3$ and $y/x = 3/4$ appear. Starting at the line $y = x$, at which the mutant and wild type phages have the same lysis time, and moving downwards, each of these ratios represents the mutant gaining the ability to carry out an additional lysis cycle. For example, the step increase in fitness observed at $y/x = 1/2$ represents the mutant finishing its second lysis cycle and beginning a third, at the same time the wild type finished its first and begins its second. Larger cell counts allow for more rounds of lysis, thus more fitness discontinuities appear as the timing of the third, fourth and fifth rounds of lysis become relevant. There would be no advantage in the mutant ensuring its fourth round of lysis completed before the wild type's third round if there are insufficient cells to support that many rounds.

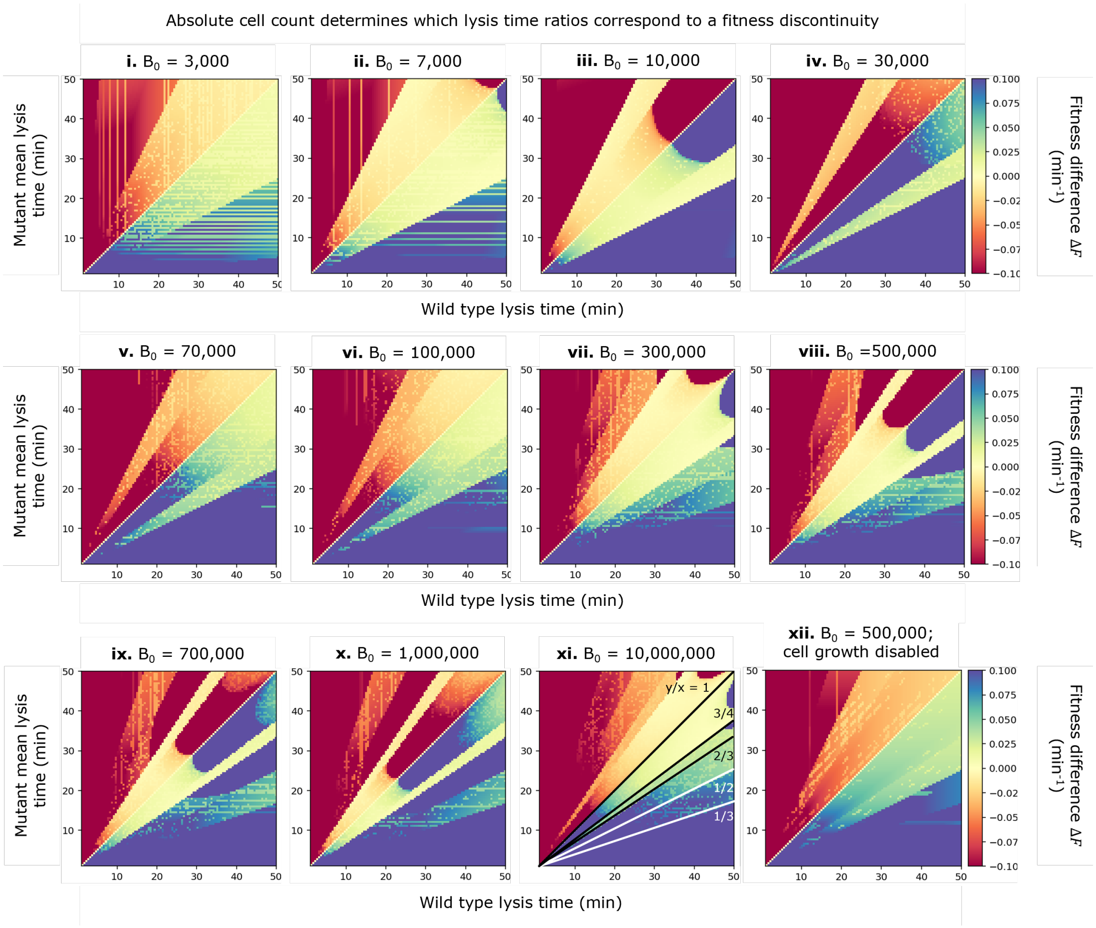


Fig. S14: Results of 'fast' simulation of a single serial passage cycle, competing a wild type phage with one mean lysis time against a mutant with another. The 'normal' model was used throughout, and intial cell count $B_0$ was varied for each panel. In all panels except xii cell growth was enabled with doubling time $T_D = 20$ minutes and no carrying capacity enforced. In panel xii, cell growth was disabled by setting $T_D = 20,000$ min.

Second, we see a screening of the fitness advantage obtained by a mutant which is only slightly faster than the wild type. In Fig S14 i, ii, vi, vii, xi, a mutant phage with lysis time one minute faster than wild type displays only a slight competitive advantage. In Fig S14 iii, iv, v, viii, ix, x, there is a region of parameter space for which such a mutant displays a significant fitness advantage (e.g. Fig S14iii, $B_0 = 10,000, \tau_{\text{wt}} = 40, \tau_{\text{mu}} = 39$). As cell count is increased, these regions seem to periodically appear at the top right corner of parameter space and move along the line $y = x$ towards the origin. These regions are first visible for initial bacteria counts $B_0$ of 7,000, 300,000 and 10,000,000, which are related by a common ratio of about 35. The significance of this value is not obvious, as it is much less than the bacteriophage's burst size of 150.

We compare Fig. S14 viii and xii, which are both initialised with 500,000 cells. In viii cells grow with a doubling time of 20 minutes, whereas in xii cell growth is disabled. This has the effect that the fitness differences in the region just below $y = x$ have been homogenised - there is no longer a region of very minor fitness difference, and region of extreme fitness difference. We take this as evidence that this region arises due to the relationship between the number of phages inoculated at $t = 0$, and the number of cells available for lysis, which is a function of both how many cells are created at $t = 0$, and how many new cells are produced by division.

#### 5.3 Implementation of fast minimal simulation

A number of control simulations were conducted using a faster, minimal simulation. This simulation makes two additional assumptions: that adsorption is instantaneous, and that lysis times and burst sizes are totally deterministic. This naturally means the minimal simulation cannot be used for investigation of lysis time distributions, but can be used to quickly measure the importance of properties such as maturation rate, eclipse period, adsorption limit, cell doubling time, and initial cell count. In the final two cases, the minimal simulation is particularly suitable as it is not object-based, so computational memory does not scale with the number of cells used, which may therefore be set arbitrarily high.

The simulation is based on eight variables, eight parameters, and three update rules, outlined in tables 1 to 4. The simulation variables are described in table 1.

**Minimal simulation variables**

| | |
|---|---|
| $t$ | the time |
| $B_T$ | the total number of bacteria |
| $B_H$ | the number of healthy / uninfected bacteria |
| $I_w$ | the number of bacteria infected by the wild type phage |
| $I_m$ | the number of bacteria infected by the mutant phage |
| $V_w$ | the number of free wild type phage |
| $V_m$ | the number of free mutant phage |
| $a$ | the total number of phages adsorbed to all living cells |

Table 1: Variables used in fast minimal simulation

The simulation parameters are described in table 2.

**Minimal simulation parameters**

| | |
|---|---|
| $B_0$ | initial bacteria count [-] |
| $T_D$ | bacterial doubling time [min] |
| $A_{\max}$ | maximum adsorbed virions per bacterium [-] |
| $\tau_w$ | wild type lysis time [min] |
| $\beta_w$ | wild type burst size [-] |
| $\tau_m$ | mutant lysis time [min] |
| $\beta_m$ | mutant burst size [-] |
| $V_0$ | inital number of each virion type [-] |

Table 2: Parameters used in fast minimal simulation

The update rules are 'adsorb', 'burst' and 'round off', implemented as follows.

**Adsorb** corresponds to as many virions as possible of one type (m or w) binding to bacteria. First, quantity $P_0$ is calculated, corresponding to the probability that a cell has no virions adsorb to it.

$$P_0 = \frac{e^{-V_w/B_T}}{B_T}$$

(i.e the $k = 0$ term of a Poisson distribution with mean $V_w/B_T$). Next, the number of virions that adsorb $\Delta V$ is taken as

$$\Delta V_i = \max(B_T \cdot A_{\max} - a, V_i).$$

That is, virions adsorb until either there are no free virions remaining, or the available bacteria are saturated.

The variables are updated as described in table 3.

| **'Adsorb' update rule** | |
|---|---|
| $t \rightarrow t$ | this step is assumed to be instantaneous, so the 'time' variable is not updated. |
| $B_T \rightarrow B_T$ | the total number of bacteria does not change. |
| $B_H \rightarrow B_H \cdot P_0$ | fraction $P_0$ of bacteria are not adsorbed to. |
| $I_i \rightarrow I_i + B_H(1-P_0)$ | healthy cells have probability $(1-P_0)$ to become infected by the appropriate phage $i \in \{w, m\}$. |
| $V_i \rightarrow V_i - \Delta V_i$ | the number of free phage is reduced by the number that were able to adsorb. |
| $a \rightarrow a + \Delta V$ | the total adsorbed phages is updated to include those just adsorbed |

Table 3: 'Adsorption' step variable update rules

**Burst** corresponds to an interval of time $\Delta t$ passing, followed by all cells infected by one phage type (m or w) lysing simultaneously, releasing their progeny. First, cell growth factor $G$ is calculated as

$$G = e^{\frac{\ln(2)\cdot\Delta t}{T_D}}.$$

The variables are then updated as described in table 4.

| **'Burst' update rule** | |
|---|---|
| $t \rightarrow t + \Delta t$ | $t$ is advanced by the time interval. |
| $B_T \rightarrow B_T - I_i + B_H \cdot G$ | the total number of bacteria is reduced by infected cells lysing, but increased by healthy cells dividing. |
| $B_H \rightarrow B_H \cdot G$ | the number of healthy cells is increased by the growth factor. |
| $I_i \rightarrow 0$ | all cells infected by the appropriate phage are lysed $i \in \{w, m\}$. |
| $V_i \rightarrow V_i + I_i \cdot \beta_i$ | the number of free phage is increased by the number of lysing cells multiplied by the burst size |
| $a \rightarrow a \cdot (1 - \frac{I_i}{I_i + I_j})$ | the total count of phages adsorbed to living cells is reduced by the fraction of infected cells which just lysed. $i \in \{w, m\}$, $j \in \{w, m\}$, $j \neq i$. |

Table 4: 'Burst' step variable update rules

**Round off** simply rounds all variables other than $t$ to the nearest integer.

The simulation begins with an an **adsorb** update called for both phage species. After that, the simulation enters a loop of checking when the next lysis event should be (which is the next smallest integer multiple of either $\tau_w$ or $\tau_m$), jumping forwards in time to that event ($\Delta t = t_{\text{nextlysis}} - t$), then calling the **burst, adsorb** and **round off** update rules. This process continues until either (a) all cells are dead, or (b) there are no infected cells or free phage remaining, which can happen if both burst sizes are 0.

The fitness difference $\Delta F$ is then determined through

$$\Delta F = \frac{\ln\left(\frac{V_m}{V_w}\right)}{t}$$

for final values of $V_m$, $V_w$, and $t$.

In order that the simulation can always uniquely determine which lysis event should happen next, the lysis times for the mutant phage are all increased by $2^{-10}$ minutes.

Given the simplicity of this simulation, we see remarkable qualitative agreement to the full stochastic simulation, including correct prediction of the existence and position of "stripes" (compare Fig 4biii leftmost column to Fig S14iii).

### 6 Assay comparison reveals weaker selective pressure in plaque assay and broadening of resonance stripes at high lysis time std

To aid comparison between the serial passage (Fig 5) and plaque expansion (Fig 6), we directly subtract the fitness advantage displayed by the mutant in one assay from another (Fig S15).

Note in particular the large blue region in the first row under the 'linear with eclipse' model, and corresponding brown region in the third row. These represent the mutant achieving fixation by population resonance in the serial passage experiment (i) but not in the plaque expansion (iii).

We use a new colour-map for this plot, as blue and brown correspond to the mutant displaying a larger fitness in one assay versus another, not displaying a fitness advantage over the wild type.

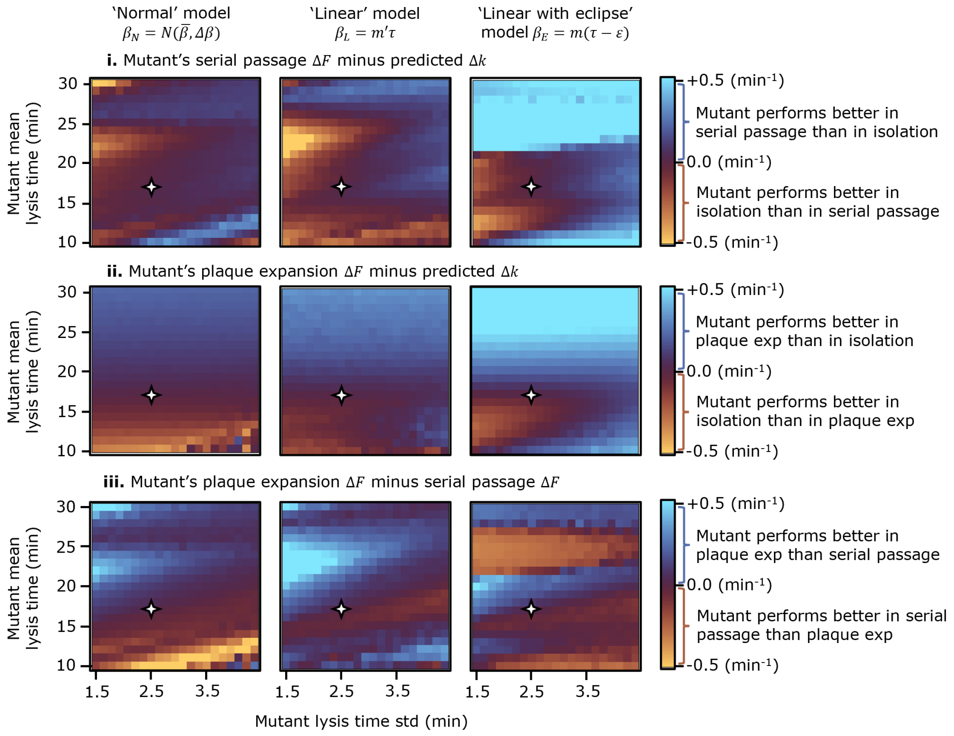


Fig. S15: Assay comparison for serial passage competition experiment, competing a wild type phage with lysis time mean 17 min, std 2.5 min against a mutant with varied lysis time mean (y axis) and std (x axis). **i.** Difference between mutant's fitness advantage in serial passage vs predicted advantage based on growth rate in isolation $\Delta F_{\mathrm{SP}} - (k_{\mathrm{mu}} - k_{\mathrm{wt}})$. **ii.** Difference between mutant's fitness advantage in plaque expansion vs predicted advantage based on growth rate in isolation $\Delta F_{\mathrm{PE}} - (k_{\mathrm{mu}} - k_{\mathrm{wt}})$. **iii.** Difference between mutant's fitness advantage in serial passage vs fitness advantage in serial passage $\Delta F_{\mathrm{PE}} - \Delta F_{\mathrm{SP}}$. Axes are scaled such that each pixel step in the y direction corresponds to a 1 minute change in mean lysis time, while each pixel step in the x direction corresponds to an equivalent fractional change in lysis time std versus the wild type value: 2.5/17 minutes. The four-point star on all panels labels the point in parameter space at which both phages have exactly the same parameters.

### 7 The secondary fitness maximum observed under the 'linear with eclipse' model is explained by population resonance

To demonstrate that population resonance is responsible for the high fitness region towards the top of the 'linear with eclipse' fitness map in Fig 5ii, we repeat the simulation using either different initial cell counts (Fig S16 i to v), or under 'conservative' conditions (no super-adsorption or decay, Fig S16vi). The initial virion titre is held constant at 100 virions of each type across all simulations. The simulation volume is co-scaled with cell count.

As the cell count is increased from 3,000 to 70,000, the high fitness region appears and disappears again, hence the name "resonance", while the high fitness region towards the bottom of the plot remains largely conserved. Note the distinction between this investigation and Fig 4.1. Here, the initial cell count, volume and carrying capacity are all rescaled by the same factor, while the initial bacteriophage titre is held constant. In Fig 4.1, the initial virion count is also rescaled by the same factor. Population resonance is highly dependent on initial MOI, but independent of absolute cell count and virion number.

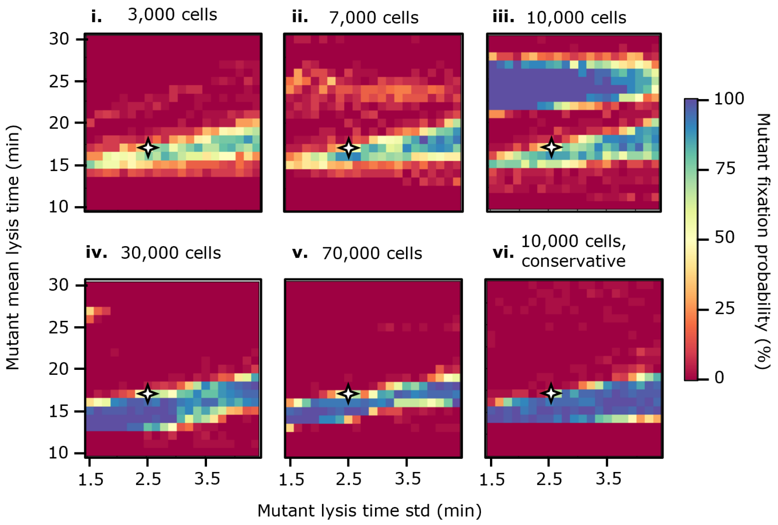


Fig. S16: This figure is a direct comparison to Fig 5. Mutant fixation probability in serial passage competition experiment, competing a wild type phage with lysis time mean 17 min, std 2.5 min against a mutant with varied lysis time mean (y axis) and std (x axis). Super-adsorption and decay are permitted in panels i to v, but disabled in panel vi. The initial number of cells $B_0$ used in each simulation can be seen above each panel. Panel iii corresponds to the set up used in Fig 5 of the main paper.

#### References


[1] Sherin Kannoly, Abhyudai Singh, and John J. Dennehy. "An Optimal Lysis Time Maximizes Bacteriophage Fitness in Quasi-Continuous Culture". In: *mBio* 13.3 (Apr. 25, 2022). Publisher: American Society for Microbiology, e03593–21. DOI: 10.1128/mbio.03593-21. URL: https://journals.asm.org/doi/full/10.1128/mbio.03593-21 (visited on 06/30/2025).

[2] Richard H. Heineman and James J. Bull. "Testing optimality with experimental evolution: lysis time in a bacteriophage". In: *Evolution; International Journal of Organic Evolution* 61.7 (July 2007), pp. 1695–1709. ISSN: 0014-3820. DOI: 10.1111/j.1558-5646.2007.00132.x.

[3] Hai Xu et al. "Biological Characterization and Evolution of Bacteriophage T7-holin During the Serial Passage Process". In: *Frontiers in Microbiology* 12 (2021), p. 705310. ISSN: 1664-302X. DOI: 10.3389/fmicb.2021.705310.

[4] Marian Dominguez-Mirazo et al. "Accounting for cellular-level variation in lysis: implications for virus–host dynamics". In: *mBio* 0.0 (July 19, 2024). Publisher: American Society for Microbiology, e01376–24. DOI: 10.1128/mbio.01376-24. URL: https://journals.asm.org/doi/10.1128/mbio.01376-24 (visited on 08/01/2024).
